# Automated Lifespan Determination Across *Caenorhabditis* Strains and Species Reveals Assay-Specific Effects of Chemical Interventions

**DOI:** 10.1101/757302

**Authors:** Stephen A. Banse, Mark Lucanic, Christine A. Sedore, Anna L. Coleman-Hulbert, W. Todd Plummer, Esteban Chen, Jason L. Kish, David Hall, Brian Onken, Michael P. Presley, E. Grace Jones, Benjamin W. Blue, Theo Garrett, Mark Abbott, Jian Xue, Suzhen Guo, Erik Johnson, Anna C. Foulger, Manish Chamoli, Ron Falkowski, Ilija Melentijevic, Girish Harinath, Phu Huynh, Shobhna Patel, Daniel Edgar, Cody M. Jarrett, Max Guo, Pankaj Kapahi, Gordon J. Lithgow, Monica Driscoll, Patrick C. Phillips

**Affiliations:** Institute of Ecology and Evolution, University of Oregon, Eugene, OR 97403 USA; The Buck Institute for Research on Aging, Novato, CA 94945, USA; Rutgers University, Dept. of Molecular Biology and Biochemistry, Nelson Biological Laboratories, Piscataway, NJ 08854, USA; Division of Aging Biology, National Institute on Aging, Bethesda, MD, 20892-9205, USA

**Keywords:** Lifespan Machine, *Caenorhabditis elegans*, automation, Thioflavin T, lifespan, CITP

## Abstract

The goal of the *Caenorhabditis* Intervention Testing Program is to identify robust and reproducible pro-longevity interventions that are efficacious across genetically diverse cohorts in the *Caenorhabditis* genus. The project design features multiple experimental replicates collected by three different laboratories. Our initial effort employed fully manual survival assays. With an interest in increasing throughput, we explored automation with flatbed scanner-based Automated Lifespan Machines (ALMs). We used ALMs to measure survivorship of 22 *Caenorhabditis* strains spanning three species. Additionally, we tested five chemicals that we previously found extended lifespan in manual assays. Overall, we found similar sources of variation among trials for the ALM and our previous manual assays, verifying reproducibility of outcome. Survival assessment was generally consistent between the manual and the ALM assays, although we did observe radically contrasting results for certain compound interventions. We found that particular lifespan outcome differences could be attributed to protocol elements such as enhanced light exposure of specific compounds in the ALM, underscoring that differences in technical details can influence outcomes and therefore interpretation. Overall, we demonstrate that the ALMs effectively reproduce a large, conventionally scored dataset from a diverse test set, independently validating ALMs as a robust and reproducible approach towards aging-intervention screening.

## Introduction

Interventions that can promote healthy human aging are likely to emerge from experimental studies performed with model organisms. Notably, metformin and rapamycin, current prominent candidate interventions for the betterment of human aging, were both first observed to extend the lifespan of laboratory animals (Harrison *et al.* 2009; Onken and Driscoll 2010). The microscopic *C. elegans* has been a widely used model in aging research for the past 30 years due to its simple genetics and its short 15-20 day lifespan. Extensive work in the field is devoted to testing compounds for the ability to extend *C. elegans* lifespan and healthspan (Castillo-Quan *et al.* 2015; Maglioni *et al.* 2016).

By their very nature, experiments that test compounds for impact on lifespan and/or healthspan are time-consuming and expensive, even in model organisms. Moreover, inconsistencies regarding the effects of some compounds on lifespan, even in relatively simple experimental systems, have been reported in the literature (Lithgow *et al.* 2017). Given the phenotypic variability that typically accompanies aging, definitive studies need to include sufficiently large cohort sizes and must be rigorously supported by replication (Lucanic *et al.* 2017; Petrascheck and Miller 2017). We previously demonstrated that robust and reproducible effects of chemical compounds on lifespan can be obtained across different laboratories using strains and species of the nematode genus *Caenorhabditis* (Lucanic *et al.* 2017). Our study also documented unexplained, but reproducible sources of variability for *Caenorhabditis* lifespan, particularly between experiments, even under a highly standardized and extensively documented series of protocols.

Variability among experiments demands multiple technical and biological replicates be performed. At the same time, however, standard survivorship assays are labor-intensive endeavors that require researchers to monitor and maintain large cohorts of animals and spend significant time microscopically assessing animal health and lifespan. Automation of survivorship assays represents a promising option for increasing throughput and enhancing reproducibility. Automated *C. elegans* lifespan assessment was introduced in 2013 with the publication of a successive image capture technology for animals on plates, based on images recorded by high resolution document scanners (Stroustrup *et al.* 2013). The table-top scanners, termed Automated Lifespan Machines (ALMs), collect images of synchronously aging cohorts that are cultured on agar plates housed on the scanner bed, at regular time intervals. Computational comparison of successive images then allows for the location of each animal to be assessed between timepoints; “non-motile” animals that consistently identify at the same location in successive images (no pixel change in animals detected between images) are scored as dead. This retrospective approach (tracking the final trajectory of an individual to a non-motile state) is much easier for the analysis of mixed populations than a prospective approach (continuous tracking of specific individuals over a lifetime), making the automated approach particularly well suited for large-scale longevity assays.

The original description of the ALM technology included comparison of Automated Lifespan Machine results with standard lifespan assays (Stroustrup *et al.* 2013). This comparison, along with our own preliminary results with the platform (Lucanic *et al.* 2016), suggested that pursuing this technology could increase throughput and perhaps reduce variability among experiments. Here we present an assessment of the ALM platform and ask whether ALM technology can be utilized to measure survivorship of different strains and species. We also tested whether the effects of chemical compounds previously shown to extend lifespan in manual assays (Lucanic *et al.* 2017) would be similarly reported by the ALM assay.

Our comparison of the ALM data we document here (from 31,983 individuals) with our previously published, manual assay data (71,252 individuals) indicated that, in general, animals are slightly shorter-lived as measured by the ALM assay, but that the rank order of strain-specific lifespan differences is generally consistent; strains that are long-lived in manual assays are also long-lived in the ALM assay. Most compounds that extended lifespan in the manual assays also extended lifespan in ALMs. However, two compounds failed to extend lifespan on the ALMs, including thioflavin T (ThT), which is highly robust and reproducible in manual assays (Alavez *et al.* 2011; Lucanic *et al.* 2017). We found ThT to be photo-sensitive, such that modification of the ALM with the addition of a blue light filter restored the beneficial effects of ThT. In fact, scanner light exposure was associated with mild negative impact on late age viability in intervention trials. Another treatment, α-ketoglutarate (AKG), induced a different outcome on ALMs vs. standard plate assays for reasons we were not able to ascertain. We conclude that the ALM platform can be utilized in large scale studies of chemical interventions in aging across genetically divergent strains, while noting that blue light filtering may be generally advised to bring ALM data in line with manual data.

## Results

To compare *Caenorhabditis* survivorship results from manual assays with results obtained from ALMs, we utilized a study design similar to that employed in our published manual assays, which comprised 22 strains of *Caenorhabditis* spanning three species (Lucanic *et al.* 2017). For the ALM study (automated analysis), each strain was setup for survival analysis using 3 independent trials, with each cohort containing over 100 animals (3 biological replicates, with each biological replicate spread over three technical replicates of 35-50 animals). This design was similar to that which we used for the manually collected data set and, as with the previous study, the data set was also produced independently in three laboratories at different geographical locations. Ideally, these experiments should result in approximately 1,080 animal observations per strain, each spread over 27 plate populations from 9 independent cohorts. The complete set contained 22 strains, yielding an approximate upper limit total of 23,760 observations. That calculation represents an upper limit since in actual survivorship studies individual animals or even whole plates of animals are lost or censored from analysis (see Discussion). The sum of final animals included for ALM data in this segment of the analysis was 13,498.

We established synchronously aging cohorts from frozen stocks of a single origin, which were continuously cultured for at least three generations post-thaw under a standard protocol at 20 °C at all three sites (<u>doi:10.1038/protex.2016.086</u>). We transferred the cohorts to agar plates arrayed on the scanners at day 7 of life (day 5 of adult life) and maintained synchrony using FuDR to prevent progeny production. The scanners captured images of each plate every 60 minutes. We called animal death based on lack of movement between successive images using the computer algorithm written for this purpose (Stroustrup *et al.* 2013). At the end of a survival study, we carefully curated data to assess the accuracy of the age-at-death calls, using a process called “storyboarding” (Stroustrup *et al.* 2013). Storyboarding involved reviewing all generated data to audit software results, verifying visually that the software made accurate estimates for times of death for each plate.

### ALM technology compares favorably to manual studies, with some differences noted

Overall, most ALM studies closely approximated outcomes of manual studies, although we noted some differences. Comparison of the survivorship between the assays indicates that the ALM populations lived frequently shorter (species average of median lifespans in days, *C. elegans* =16.8, *C. briggsae*= 20.6, *C. tropicalis*=18.0) than did the manually scored populations (species average of median lifespans in days, *C. elegans* =19.5, *C. briggsae*= 23.6, *C. tropicalis*=22.8), with an overall ~14% shorter mean lifespan (Figure 1A and Online Resource 1). We suspect that the shortened lifespans on ALMs may be related to the frequent exposure to intense light, which includes wavelengths in the visible range that can shorten *C. elegans* lifespan (De Magalhaes Filho *et al.* 2018).

**Fig. 1.**
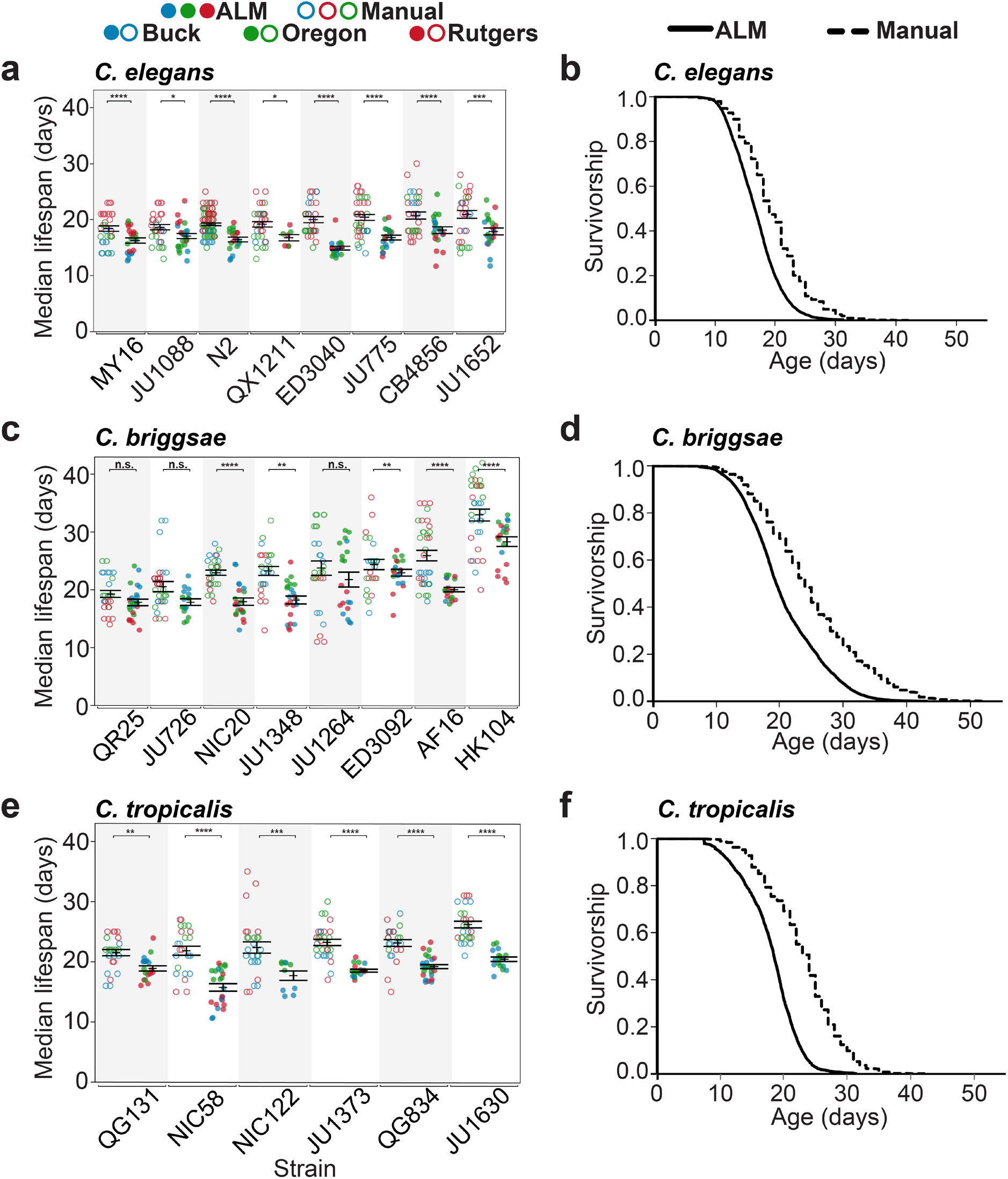
Survival differences among *C. elegans*, *C. briggsae* and *C. tropicalis* species are reported similarly by manual and automated lifespan analysis. (a,c,e) Comparison of lifespans from manual and ALM survival analyses. Median lifespans from eight *C. elegans* (a), eight *C. briggsae* (c) and six *C. tropicalis* (e) strains. Strains are ordered from shortest to longest average median lifespan in the manual assays. Each point represents the median lifespan from an individual plate trial conducted at one of the three CITP sites (Blue-Buck Institute, Green-Oregon and Red-Rutgers); all sites contributed three replicates of three plates tested/trial; The bars represent the mean +/- the standard error of the mean. For the manual data (open circles) ~35 animals were used to initiate each plate; For the ALM data (closed circles) ~50 animals were used to initiate each plate to account for worm loss in initial transfers to give a similar yield as in the manual assays. Asterisks represent *p*-values (\*\*\*\**p*<0.0001, *** *p*<0.001, ** *p*<0.01 and * *p*<0.05) from the CPH model when comparing the lifespans measured by ALM versus manual assay. Summaries of the parent data used to generate these graphs are included in Online Resource 1. (b,d,f) Survivorship curves from data of combined trials for all *C. elegans* (b), *C. briggsae* (d) and *C. tropicalis* (f) strains. Manual data are from Lucanic et al (2017).

The relative orders of mean lifespans we recorded were generally similar between the manual and automated platforms (Figure 1A-C, Online Resources 1 and 2). The ALM-reported mean lifespans fall within a relatively narrow range for the *C. elegans* strains (ED3040=15.2 to CB4856=17.9 days). In addition to having the shortest mean lifespan on the ALMs among the *C. elegans* strains, ED3040 also exhibited the largest decrease among the *C. elegans* strains in median lifespan relative to manual assays (5.2 days), suggesting a modestly enhanced ALM stress for this strain. *C. tropicalis* strain JU1360 and *C. briggsae* strain AF16 were the exceptional strains that exhibited the biggest difference in mean lifespan between the two datasets and shifted their relative positions in survival comparisons. The *C. tropicalis* strains showed (on average) the largest differences between the manual and ALM datasets when compared to the other two species. This difference is not the result of how missing worms are scored (“censoring” Online Resources 3-5), which could result from a species-specific sensitivity to lab conditions.

Overall, we recorded survival data from 22 strains with 9 repeat trials each spread equally over three geographical sites. As was the case for our manual survival assays (Lucanic *et al.* 2017), we measured some variability across the ALM dataset, and we statistically evaluated the sources of this variation. We compared the sources of this variability to those in the manual dataset by identical statistical methods, partitioning variation among potential sources of error using a general linear model (GL) (Table 1 and Online Resource 6). We observed similar variability arising from genetic differences, reproducibility among labs, and reproducibility within labs, for ALM vs. manual comparisons. For manual assays ~19.7% of the total variance is attributed to genetic sources, ~7.5% of the total variance is attributed to overall lab-specific effects, and ~15.4% of the total variance is attributed to variation within labs (Lucanic *et al.* 2017). For ALM assays, sources of variation were extremely similar, with ~21.7% of the variation attributed to genetic differences, ~9.4% variation attributed to overall lab-specific effects, and ~10.5% attributed to variation within labs. None of the patterns of variation between the experimental approaches is notably different except for the among-trial variation. The ALM data exhibited lower among-trial variation within labs (9.3% manual; 2.5% ALM) that was not linked to a reduction in the variation among individual culture plates.

**Table 1.** Comparison of reproducibility of longevity estimates from manual and automated lifespan assays within and between labs for the baseline analysis of 22 strains across three species with no added compounds. Results for manual assays are from Lucanic et al. (2017).

| Source of Variation | Manual Assays | Automated Assays |
| --- | --- | --- |
| **Genetic Variation** | **19.7** | **21.7** |
| Among species | 11.7 | 10.4 |
| Among strains w/in species | 8.0 | 11.3 |
| **Reproducibility Among Labs** | **7.5** | **9.4** |
| Among labs | 0.0 | 1.9 |
| Lab x species | 0.6 | 0.6 |
| Lab x strain | 6.9 | 6.9 |
| **Reproducibility Within Labs** | **15.4** | **10.5** |
| Among experimenters or scanners | 0.0 | 0.0 |
| Among trials w/in lab | 9.3 | 2.5 |
| Among plates w/in trials | 6.1 | 8 |
| **Individual variation** | **57.4** | **58.3** |
| **Total** | **100.0** | **100.0** |
| Total number of observations | 26,333 | 13,498 |

Confidence intervals were only computable for the manual assays.

### Comparison of bi-modal aging phenotypes

Most of the reduction in among-trial variation in the ALM assays appears to have been generated by more consistent strain-specific responses across automated assays. We previously showed that for some strains, the variability of survival among cohorts is not normally distributed, such that replicate cohorts were either short-lived or long-lived (we refer to this as bi-modal survival; Figure 2, top graphs; (Lucanic *et al.* 2017)). Bimodal survival had not been reported previously for nematodes, although there is now evidence that hermaphrodites can exhibit greatly reduced lifespans in the presence of males (Maures *et al.* 2014; Shi and Murphy 2014). Males are absent from our experimental populations. The mechanism at play for observations is not clearly understood, although we have suggested that bimodal survival suggests that populations can adopt alternative physiological states. Since the manual and ALM methods feature differences in nematode handling (for example, the ALM assay requires much less technician manipulation such as repeated animal transfer to fresh plates but includes repetitive light exposure), we were curious whether cohorts analyzed by the ALMs would exhibit the bi-modal survival outcomes. In other words, comparison of outcomes from manual and ALM technologies could indicate whether bimodal survival traits are induced by experimental manipulation aspects of the survival studies vs. being inherent to the natural biology of the animals.

**Fig. 2.**
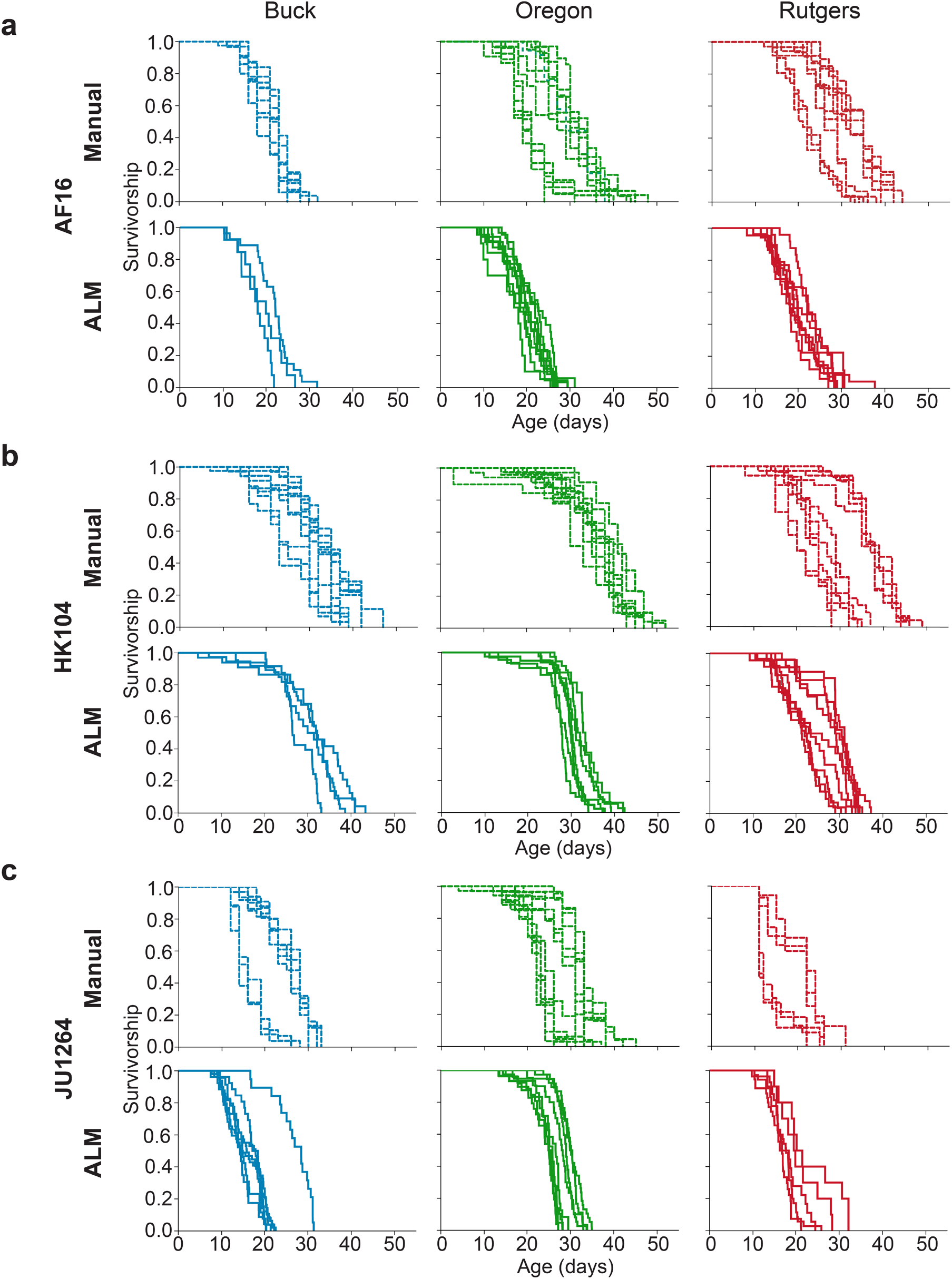
ALM lifespan analysis does not exhibit the strong bimodal distribution of lifespan curves observed in manual assays. Survivorship curves from *C. briggsae* strains AF16 (a), HK104 (b), and JU1264 (c) are presented. Data were generated by manual lifespan and ALM analysis at all three CITP sites. The qualitatively bimodality of survivorship curves observed in manual analysis in at least two of the three independent sites for each of the *C. briggsae* strains does not stand out in the ALM data set. Manual data are from Lucanic et al (2017).

Our previous work revealed that *C. briggsae* strains exhibited bi-model survival and that these outcomes were apparent in at least two of the participating *Caenorhabditis* Intervention Testing Program (CITP) labs for all strains. In the ALM assays, there is some indication that the bi-model pattern persists occasionally in some strains (e.g., HK104 at Rutgers and JU1264 at all three sites; Figure 2), but in general the differences are slight and might not have been noticed in the absence of the *a priori* expectation built upon the manual assay results. This difference manifests itself statistically as a four-fold decrease in among-trail variation for the ALM assays. We infer that assay conditions can influence the bimodal longevity feature of some strains.

### Comparison of compound treatment effects on lifespan between manual and ALM platforms

We next examined whether use of the ALM platform would return chemical intervention results similar to our conventional manual dataset. To address this question, we again set up an experimental scheme that would yield a dataset that could be compared to our previous study (Lucanic *et al.* 2017). We selected five chemical treatments that we had previously characterized using CITP experimental design: NP1, resveratrol, propyl gallate, AKG and ThT. NP1 is a compound that exerts positive effects in both *C. elegans* and *C. briggsae* strains that appears to engage dietary restriction-like metabolism (Lucanic *et al.* 2016); resveratrol is well known as a positive bioactive component of red wine (Wood *et al.* 2004); propyl gallate (Wood *et al.* 2004) and AKG (Chin *et al.* 2014; Mishur *et al.* 2016) were known to exert potent lifespan extension in *C. elegans* N2; and ThT is an amyloid binding compound that extends *Caenorhabditis* lifespan (Alavez *et al.* 2011; Lucanic *et al.* 2017). We used these five chemical treatments and two solvent controls (aqueous or DMSO based) to test for longevity responses across six different strains of two species (*C. elegans* and *C. briggsae*; Figures 3 and 4, and Online Resources 7-10). Our ALM protocol featured manual transfer of adult animals onto the surface of agar plates that contained food and either compounds or vehicle control. On the fifth day of adulthood (approximately 20% of the lifespan of the wild type laboratory strain *C. elegans* N2), we transferred animals to fresh scanner plates +/- compound, positioned the plates on the scanner bed, and initiated automated image capture. We scanned plates every hour until all animals were counted as dead. We then processed images using software developed by Stroustrup et al. (2015) (https://github.com/nstroustrup/lifespan), with human quality control posthoc examination of videos to verify that software death calls were accurately timed. We estimated that on average the ALM captured greater than 50% of the possible deaths under the experimental conditions (Online Resources 3 and 9).

**Fig. 3.**
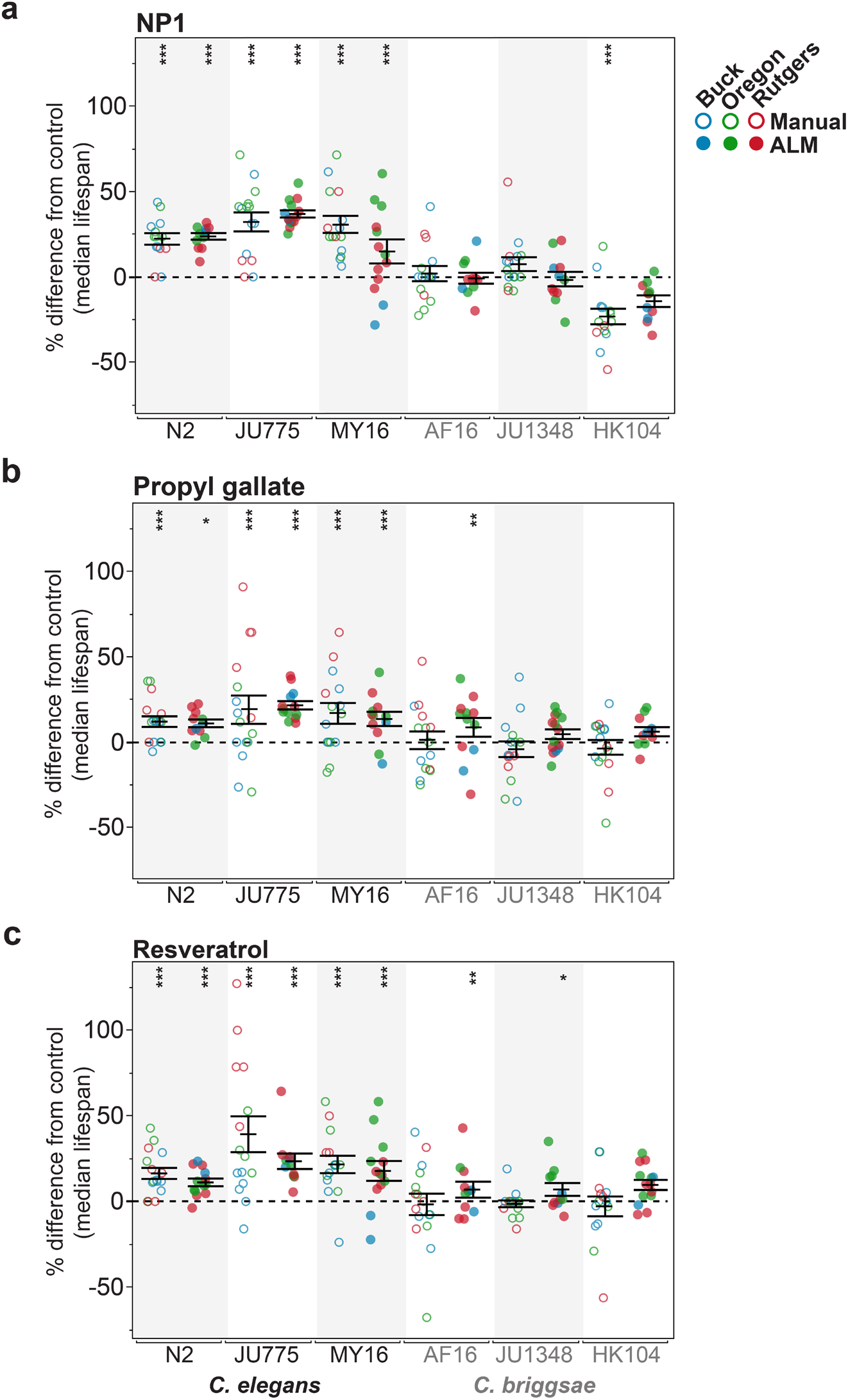
Several lifespan compound interventions are reported similarly by manual and ALM lifespan analysis. The change in median lifespan under adult exposure to NP1 (a), propyl gallate (b) or resveratrol (c) are shown for three *C. elegans* (N2, JU775, and MY16) and *C. briggsae* (AF16, JU1348, and HK104) strains. Each point represents the change in median lifespan from an individual plate trial relative to the specific control conducted. The bars represent the mean +/- the standard error of the mean. Replicates were generated at the three CITP sites (Blue-Buck Institute, Green-Oregon and Red-Rutgers). Lifespans were measured by both manual (open circles) and ALM (closed circles) survival analyses. Asterisks represent *p*-values (\*\*\*\**p*<0.0001, *** *p*<0.001, ** *p*<0.01 and * *p*<0.05) from the CPH model when comparing the lifespans under compound exposure versus the lifespans exposed to the vehicle control. Summaries of the parent data used to generate these graphs are included in Online Resource 10.

**Fig. 4.**
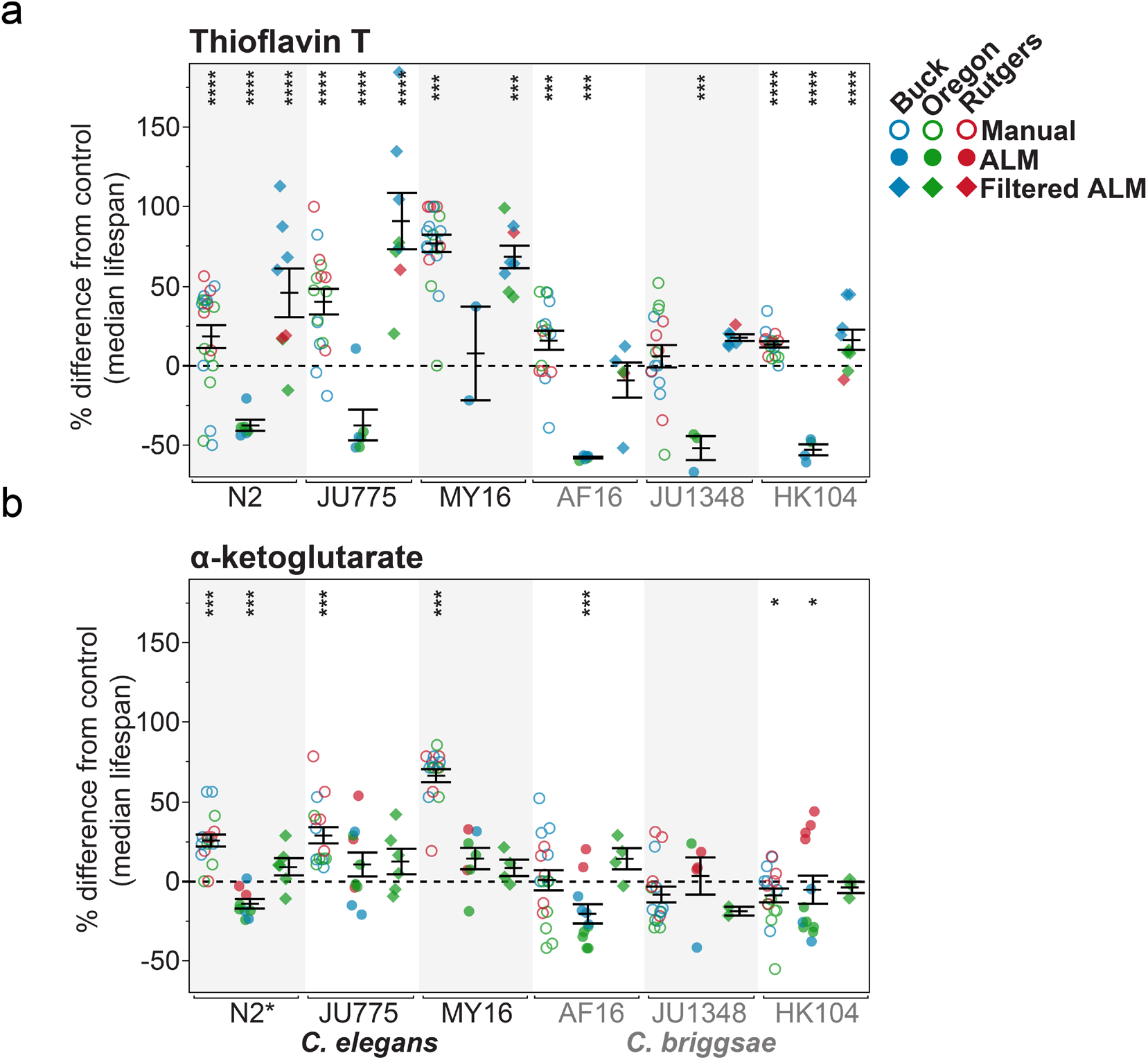
Thioflavin T lifespan effects are reversed by light exposure in ALM analysis. Changes in median lifespan under adult exposure to thioflavin T (a) or α-ketoglutarate (b) are presented. Each point represents the percent difference in median lifespan from an individual plate trial relative to the control. The bars represent the mean +/- the standard error of the mean. Lifespans were measured by both manual (open circles) and ALM (closed circles) survival analyses. (a) Median lifespans were extended to variable degrees in all strains under thioflavin T exposure when measured using manual analysis, while ALM analysis gave consistently shorter median lifespans. Automated analysis with filtered light (closed diamonds) restored lifespan extension of thioflavin T for all strains except AF16 and JU1348 where filtering eliminated the reduction in lifespan from thioflavin T exposure while not reaching statistical significance for lifespan extension. (b) α-ketoglutarate extended median lifespan in *C. elegans* strains in manual lifespan analyses, while no lifespan extension was observed in ALM analysis. Filtering the light during ALM analysis did not restore lifespan extension under α-ketoglutarate exposure. Asterisks represent *p*-values (\*\*\*\**p*<0.0001, *** *p*<0.001, ** *p*<0.01 and * *p*<0.05) from the CPH model when comparing the lifespans under compound exposure versus the lifespans exposed to the vehicle control.

Our analysis of the ALM chemical treatment data set indicated that results from three compounds NP1, resveratrol, and propyl gallate, were similar to what we had previously reported for the manual assays (Figure 3, Online Resources 7, 9-11). Specifically, we find that NP1 is effective across *C. elegans* strains but less so in *C. briggsae* as determined by both manual and ALM approaches. Likewise, resveratrol and propyl gallate confer a modest median lifespan extension in *C. elegans* but are less potent (if effective at all) in *C. briggsae* strains. These data support that the ALMs can recapitulate outcomes obtained using manual assays for compound effects. Since there are considerable savings in technician labor for the ALMs, ALM use represents an opportunity for enhancing bandwidth for compound intervention lifespan experiments.

On the other hand, we did note significant compound-specific differences from our manual derived dataset for ThT and AKG (Figure 4, compare open and filled circles, Online Resource 12 and 13): ThT and AKG were strikingly effective at promoting longevity in our manually-derived dataset but exhibited the opposite effect in the ALM assays--shortening lifespan in all but one strain, MY16, where they had no effect. For example, ThT treatment of each of the other five test strains *shortened* the median lifespan so substantially that this compound would be considered toxic in the ALM environment. Comparison of manual vs. ALM approaches thus underscores the impact of distinctive methods of lifespan determination for a subset of compounds.

### Filtering blue/UV light restores the positive effects of thioflavin T on lifespan

We considered why, for some compounds, contrasting results were observed for the ALMs and the manual assays. We first excluded that there were major differences in the reproducibility of longevity estimates for manual and ALM assays across the six strains and two species (Table 2, Online Resources 14-17). We found no significant difference in variation arising from genetic differences, or reproducibility within labs. We did note small but significant differences in variation among strains between labs (manual 0.2% vs. automated 8.2%) but these differences are not sufficient to explain the dramatic changes in chemical intervention outcome we observed. Having ruled out major experimental variation in outcome, we considered the potential impact of the most fundamental difference between manual and ALM assays, differences in light exposure. In the ALMs, animals and the chemicals in the plates they are maintained on are subjected to a light scan every fifteen minutes (see materials and methods) of their lives after placement on the scanners. Blue light has been documented to shorten lifespan in standard manual plate assays (De Magalhaes Filho *et al.* 2018), and many compounds are photolabile. We noted that structural motifs in ThT could be predicted to be photolabile. Moreover, modified forms of ThT are known to be produced by oxidation (Al-Maqdi *et al.* 2017). Indeed, we previously observed that exposure to high concentrations of ThT was highly toxic (Alavez *et al.* 2011), which might be explained by toxic breakdown products. We therefore speculated that exposure to high intensity light in the ALMs might modify ThT, resulting in the production of a species that is toxic to *Caenorhabditis* nematodes. To address this possibility, we obtained the reported frequency spectrum produced by the scanners and noted a prominent blue/UV emission. We modified the ALMs with a filter that reduced the blue/UV emission and repeated the experiments with ThT. Under the modified ALM, ThT reproducibly and robustly extended lifespan (Figure 4, Online Resources 8a and 18), which is especially evident in the *C. elegans* strains. In some cases, ThT extended lifespan to a greater extent than observed in the manual assays. Thus, for the ThT intervention, changing the blue/UV light exposure of the test plates conferred a profound difference in outcome. These data both underscore the potential for different outcomes using ALMs and suggest a light filter-based approach toward eliminating complications of testing some photolabile compounds.

**Table 2.** Comparison of reproducibility of longevity estimates for manual and automated assays for pharmacological intervention for three different compounds (plus controls) for six strains across two species. Results for α-ketoglutarate and Thioflavin T are excluded from this analysis. Variance estimates for the compound trials are the averages across all compounds as estimated from a single general linear model. Estimates for manual assays are derived from Lucanic et al. (2017) for the subset of compounds used in this study.

| Source of Variation | Manual Assays | Automated Assays |
| --- | --- | --- |
| **Genetic Variation** | **44.9** | **33.1** |
| Among species | 32.5 | 12.8 |
| Among strains w/in species | 7.1 | 13.3 |
| Species x Compound | 3.2 | 5.1 |
| Strain x Compound | 2.0 | 2.0 |
| **Reproducibility Among Labs** | **0.7** | **8.3** |
| Among labs | 0.0 | 0.0 |
| Lab x species | 0.4 | 0 |
| Lab x strain | 0.2 | 8.2 |
| Lab x compound | 0.0 | 0.1 |
| **Reproducibility Within Labs** | **8.8** | **8.3** |
| Among experimenters or scanners | 1.5 | 0.0 |
| Among trials w/in lab | 2.2 | 2.8 |
| Among plates w/in trials | 5.1 | 5.5 |
| **Individual variation** | **45.7** | **50.3** |
| **Total** | **100.0** | **100.0** |
| Total number of observations | 15,683 | 8,846 |

Confidence intervals were only computable for the manual assays.

Increased sensitivity of worms to light in the presence of ThT raises the possibility that light on the ALM per se might be a source of increased mortality. Indeed, median lifespan can be increased by as much as nine days when filtered light is used instead of the full spectrum scanner light (Figure 5, Supplementary Table 6). The largest light effects appear to be in late life mortality, with maximum lifespan displaying a much larger increase than median lifespan per se, especially in *C. elegans*. We therefore consider it likely that light exposure effects are a cause of the overall decrease in lifespan using the ALM versus manual approaches highlighted (Figure 1).

**Fig. 5.**
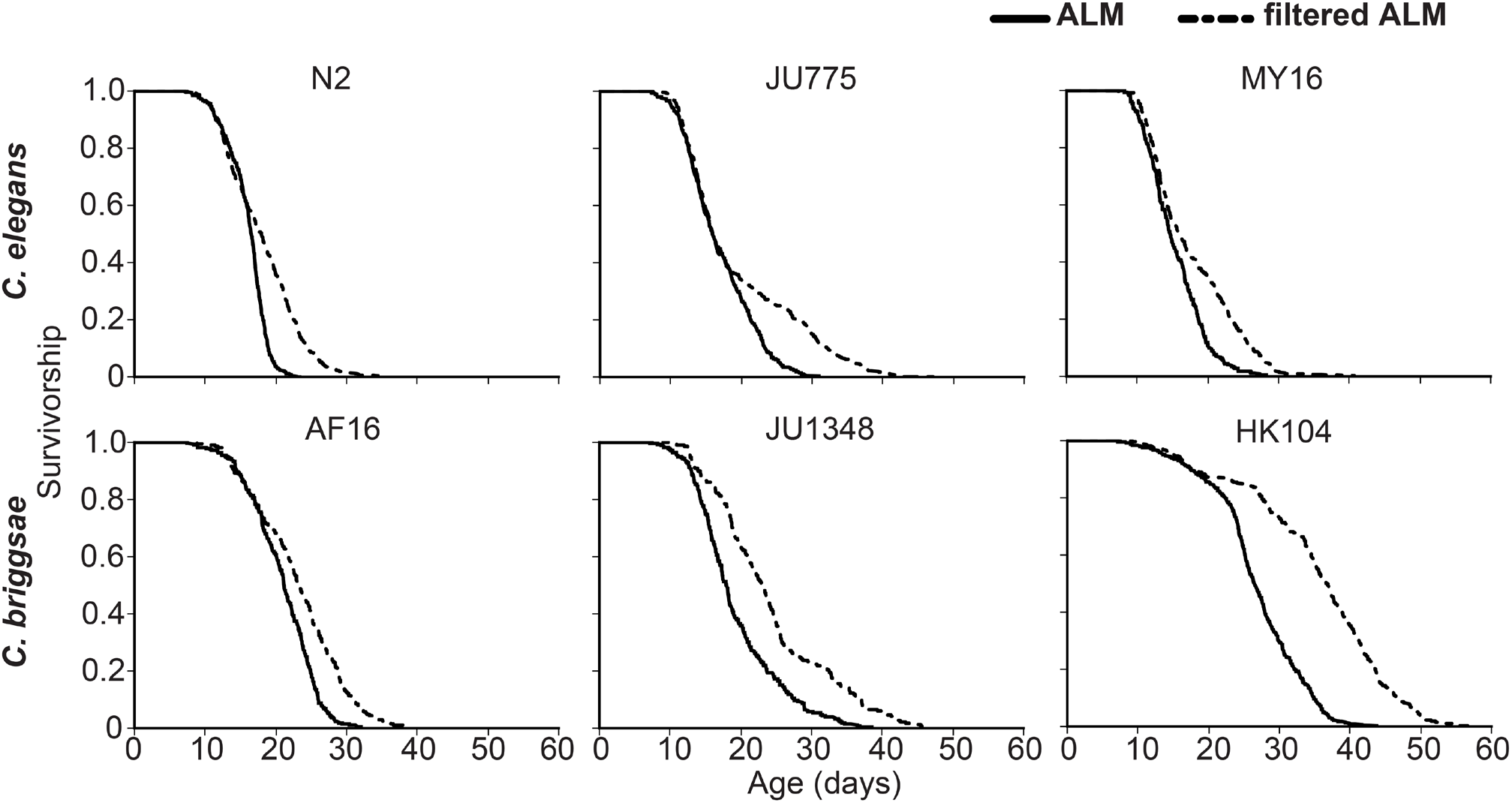
Light filtering during ALM analysis changes late life survivorship. Survivorship curves for ALM assays with (dashed line) or without (solid line) light filtering for three *C. elegans* (N2 *p*<0.001, JU775 *p*=0.004, and MY16 *p*<0.001) and three *C. briggsae* (AF16 *p*=0.051, JU1348 *p*=0.001, and HK104 *p*<0.001) strains. The *p*-values were calculated using the CPH model when comparing the lifespans measured with versus without light filtering (see Materials and Methods).

We also tested whether the same light filter approach might address the discrepancy with manual outcomes for AKG. In contrast to the results for ThT, filtered ALM experiments do not rescue the positive effects of AKG observed in manual assays (Figure 4, Supplementary Figure 4B). AKG is an acid, and our 192 mM stock solution had a measured pH of 1.56. Treatment of the buffered plates could therefore result in a lower final plate pH, while the bacteria present on the plate during the treatment would be expected to experience a significant transient drop in pH. Consistent with this, we observed greatly reduced bacterial cell viability after AKG treatment. We therefore retested AKG on the ALMs using stock solutions that had been titrated with 10M NaOH to a pH of 6 before addition to the agar plates. The pH adjustment rescued bacterial cell viability but had no impact on AKGs effect on lifespan, and thus is unlikely to be the source of the difference between methods (Online Resource 19). Thus, the mechanism of efficacy loss for AKG in the ALM tests (particularly for *C. elegans* strains) remains unexplained.

Overall, our analyses of large data sets and multiple test compounds reinforce our previous conclusions that methodology details for intervention testing can have a profound effect on experimental outcome (Lucanic *et al.* 2017). We suggest that ALM modifications that filter blue light is one strategy for improving alignment of manual and automated lifespan analyses, at the same time we underscore attention to vigilance regarding approach-specific outcomes in longevity studies.

## Discussion

The rapid rise in average lifespan over the last several decades has fueled understanding the biological basis of aging as one of the pressing issues in human health. Concomitant with this need, researchers have made unprecedented progress in both deciphering the genetic bases of pathways that enhance longevity in model organisms, and identifying candidate compounds that may increase the period of healthy aging in humans. A major challenge in the field, however, is that aging itself is a very difficult phenotype to study and, in particular, lifespan is impacted by complex genetic, environmental, and stochastic factors. The highly multifactorial nature of aging as a biological phenomenon means that there is a great deal of variation among individuals, even in highly controlled experiments against uniform genetic backgrounds. The impacts of this variation are several fold. First, a large number of individuals must be measured to get accurate estimates of survivorship over time. Second, unknown and undetected sources of variability across experiments raises the possibility that individual aging experiments will be particularly difficult to reproduce. The use of automated assays in the assessment of longevity effects have the potential to address both of these concerns, or at the very least allow for the comprehensive assessment of these factors as potential sources of error in aging research. Here, we find that the *C. elegans* Automated Lifespan Machine (ALM) generates results that are similar to standard manual lifespan assays in most cases. Indeed, we find that large data sets focused on manual and automated lifespan analysis generate similar distributions of reproducibility. At the same time, we find that increased light exposure under the ALM approach can for some tested compounds, lead to important differences, both through a generalized stress that leads to an across-the-board decrease in longevity regardless of genetic background/intervention treatment and through specific compound interactions with light.

### Reproducibility in ALM-generated longevity data

We have previously noted prominent studies that have failed to reproduce aging interventions between labs despite strenuous efforts that researchers made to define the factors that create such differences (Lithgow *et al.* 2017). Many factors have been offered to explain discrepancies, including a lack of detailed methodology, differences in reagents, differences in experimenter technique, and divergent statistical analyses, among others. The *Caenorhabditis* Intervention Testing Program (CITP) is designed to discover chemical compounds (drug-like molecules) that extend lifespan and healthspan across a genetically diverse test set. The CITP effort requires an extensive collaborative effort that maintains standardized protocols. By using distinct *Caenorhabditis* strains, we are also able to determine whether the effects of a compound are robust across genetic background. Broad impact across a genetically diverse population is important for the goal of prioritizing compounds for extended research in more complex animals, including rodents.

We considered whether use of ALMs might enhance experimental reproducibility, as automation can increase efficiency and precision. Conventional worm manual lifespan assays appear fairly simple at first examination. Aged cohorts of easily cultured *C. elegans* are scored periodically and the fraction dying in given time period (usually 24 hrs) is recorded. Training is required to differentiate an old, sick worm from a dead worm, but the protocol appears deceptively straightforward compared to more complex assays in biology. However, there are variables that may affect outcomes, including media batch differences, food (*E. coli*) growth conditions, animal transfer technique, fungal and bacterial contamination, incubator and lab temperatures, microscope stage temperature and more. Automation of nematode lifespan assays offers great potential but is subject to some of these challenges and introduces new limitations of the technology itself.

Our results demonstrate that use of the Automated Lifespan Machines allows us to capture data that are quantitatively similar to our conventional manual assays and emphasize their utility in studying the effects of chemical compounds. The data on *C. elegans* N2 reproduce findings of Stroustrup et al in which the Lifespan Machines were tested extensively with similar experiments (Stroustrup *et al.* 2013, 2016). Here we have demonstrated that in addition to *C. elegans* N2, other strains and species can also be analyzed using the ALM system. One reason that the technology recapitulates manual survival assays is that agar-based culture conditions for the lifespan machines closely match the manual assays.

### Regarding the question of increased throughput

We do note that the use of the ALM platform incorporates a “storyboarding” mechanism for manual override of machine death calls, which we used to audit machine death calls in a consistent manner. Storyboarding served mainly to remove duplicates and non-worm objects, misidentified as worms. While we initially used these to alter the exact time of death called by the machine, as we introduced strain-specific use of the posture files to increase accuracy, this generally became unnecessary as the algorithm nearly always made the correct call.

ALM use requires considerably less human scoring during a survival assay than does manual assay (manual life/death determinations every other day or so). On the other hand, we did find that storyboarding after the ALM study required a considerable amount of person-time to complete (about 1-2 hours/scanner for an experienced auditor). We have found that individuals trained for part-time work storyboarding can efficiently advance this effort. Still, with storyboarding required, the overall time devoted/survival assay provides only a modest increase in efficiency over conventional manual studies. Nevertheless, the temporal resolution of the ALM experiments is an order of magnitude greater than for manual assays, so it is not only throughput per se, but much greater resolution that is an advantage in the ALM system.

### Censoring in ALM lifespan determination

Factors in plate/animal loss can include contamination, animals leaving the culture plate, animals experiencing an egg laying failure that induces uterine defects, and accidental killing during transfer. All but the last of these events occur in both automated and manual assays. In survivorship assays, most of these non-natural death events result in an animal being censored from the study, at the time at which it is detected as lost. These censored events are then included in the data set to calculate the survivor function and so influence estimations of median lifespan and significance tests. The censored events are therefore important factors in survivorship studies and while the numbers vary, they can represent a large percent of animal observations in a data set. Analysis of the number of censored animals from our previous study ((Lucanic *et al.* 2017); Online Resource 5) indicated that 20% of the total animal observations were from censored animals. The percent of animals that were censored varied dramatically amongst strains and may be representative of their propensity to crawl off the plate, burrow under the agar, or experience a failure of egg-laying. Specifically, we observed a range from 8% censored in JU1652 to 36% censored in JU1264 (Online Resource 5). In our use of the ALM, we do not record censored animals. Only animals observed to have actually died are recorded in our ALM data set. Still, our comparisons of the manual and ALM data sets suggest that survival interpretations are similar, and thus we conclude the practice of censoring in manual studies does not introduce a major difference in interpretation of experimental outcome.

### Culture differences in manual vs. ALM

We previously demonstrated that at least one compound, ThT extended the lifespan of three distinct species in manual lifespan assays. We were therefore surprised when early experiments with the ALMs indicated that compound-treated cohorts lived shorter than the vehicle controls. There are actually many differences in protocols between the manual assay and the ALMs, a particular one of which includes exposure to compound. In the manual assay, animals are moved to plates containing fresh compound every 3-4 days. In contrast, once the worms are transferred to the ALMs at day five of adulthood, they remain on the same agar plate until the assay is complete. Since we do not know the stability of the compound in the agar, or the pharmacokinetics of the compound in the worms, we cannot predict if this difference in protocol might result in a significantly different outcome. If of sufficient interest, chemical tests can be used to address compound stability over the course of an ALM study.

### Light exposure can influence ALM outcomes, with particular potency for specific compounds

Another obvious difference between manual and ALM studies is the exposure of the agar plates to intense light every 15 mins during the scanning process. Exposure to light can be detrimental to *C. elegans* (De Magalhaes Filho *et al.* 2018) and our analysis indicates that this is true in our set of test species as well (Figure 5). Together with potential temperature fluctuations if scanners are not housed in incubators, a generally shorter lifespan in the ALMs compared to manual assays might be anticipated.

In the case of thioflavin T, we also considered whether the light influenced the compound itself. Thioflavin T is known to become modified upon oxidation, and photolability could result in disruption of the efficacious structure of the chemical or the generation of a toxic species following light exposure. To begin to address these possibilities, we modified the standard operating procedure for ALMs methods (Stroustrup *et al.* 2013) to include a blue light (~100% opacity 400-460 nM) filter. This modification resulted in lifespan extension by thioflavin T, supporting that in this case, light interaction with chemical was responsible for the difference in outcomes. The ThT studies clearly demonstrate that modest differences in protocol can result in radically different outcomes. In the case of ThT, we were able to identify the critical difference in methodology that explained differences. We have not been able to identify the cause of the AKG failure to extend lifespan on the ALMs in contrast to the robust lifespan extension we observed with manual assays (pH adjustment did not matter). The manual assays require repeated transfers of the aging worms onto fresh plates whereas once the ALM plates are on the scanners, the worms are not handled further. Thus, in manual assays, a higher lifetime exposure to AKG and lower physical stress may contribute to differences.

In general, then, the ALM light is likely to introduce an overall more stressful environment than manual assays, regardless of any compound-specific interactions with the scanning environment. In some cases, the slight stress may actually make it easier to reveal the generally protective effect of a compound intervention, as seems to be the case for propyl gallate and resveratrol in *C. briggsae* (Figure 3).

Given these light effects, why would one not want to use filters at all times? Unfortunately, the filters make the overall ALM assay less effective. Because the filters block a significant fraction of the light, the scanners compensate by increasing the scan duration. This, together with the reduced illumination, makes it more difficult for the automated software to detect the movement of individual worms, and makes it much more difficult for humans to verify each worm death during the storyboarding process. Overall, then, given the overall consistency of our results, we suggest that unless there is a clear indication that light exerts a direct impact on the efficacy of a compound, we have chosen not to use a filter under our standard ALM assay conditions. Newer models of the document scanner use LED light sources instead of fluorescent light and may have reduced light-induced stress/mortality effects. This question is currently under investigation in our labs.

## Conclusion

Longevity is a notoriously difficult phenotype to assess accurately within experimental settings. Stochastic variation among individuals tends to dominate many direct interventions aimed at extending health and lifespan. We find that the Automated Lifespan Machine is a viable addition to the effert to generate highly reproducible science within the *C. elegans* longevity community. Its greatest benefits are a tremendously increased temporal resolution and a strong reduction in labor to determine the lifestate of each individual in an experiment. Most importantly, we are able to reproduce the majority of our findings regarding strain and species-specific differences in longevity, as well as the effects of specific chemical interventions on lifespan. However, the system is not in fact fully automated, as the images must be curated by a human after the experiment, which can take a fair bit of time. It is also clear that greatly increased light exposure under the ALM approach is itself stressful to the organism. This can be especially confounding when the response to a compound itself is light dependent, as we found with Thioflavin T. So overall, ALM and manual approaches are complimentary, but each longevity assay is ultimately unique to itself and is likely to measure a somewhat different longevity profile. Such is the case with all experimental conditions in longevity assays (e.g., temperature, food type, osmolarity), and so there is not in fact any objective “natural” system for laboratory studies of longevity. In the end, each assay must be used in the context of its appropriate controls and each conclusion regarding longevity interventions must be made in the context of the environment in which they are tested.

## Materials and Methods

Data and protocols for the manual lifespan analyses were previously published ((Lucanic *et al.* 2017; Plummer *et al.* 2017)). Detailed automated lifespan analysis protocols, forms, and checklists have been made available online at https://doi.org/10.6084/m9.figshare.c.4580546. Experimental protocols in brief are as follows:

### Nematode strains and culturing

The 22 nematode strains used here are natural isolates (except N2, which has undergone domestication) of the three hermaphroditic species of *Caenorhabditis* and include eight *C. briggsae* strains (AF16, ED3092, HK104, JU1264, JU1348, JU726, NIC20 and QR25), six *C. tropicalis* strains (JU1373, JU1630, NIC122, NIC58, QG131 and QG834), and eight *C. elegans* strains (CB4856, ED3040, JU1088, JU1652, JU775, MY16, N2 and QX1211). All automated analyses used the same frozen strain stocks (see (Lucanic *et al.* 2017) for description) as the manual analyses except for the subset of the α-ketoglutarate experiments with the strain noted as N2-PD1073. N2-PD1073 is a clonal line derived from the N2 strain VC2010 that was used in the generation of a new N2 reference (“VC2010-1.0)” genome (Yoshimura *et al.* 2019). All nematode culturing was done at 20 °C. Experimental temperatures were verified using previously used temperature data loggers (Lucanic *et al.* 2017; Plummer *et al.* 2017) or custom 16-channel temperature recorders (Banse *et al.* 2019).

### Survival studies

ALM SOPs were generated by modifying the protocols described for the Automated Lifespan Machine (Stroustrup *et al.* 2013) to align them with the previously described CITP manual lifespan protocols ((Lucanic *et al.* 2017; Plummer *et al.* 2017); CITP https://doi.org/10.6084/m9.figshare.c.4580546). Specifically, worms were transferred to control or compound treated plates supplemented with 51 µM FUDR (versus 40.6 µM) starting at the first day of adulthood (versus late L4). Animals were transferred to fresh plates twice more before the plates were loaded into the Automated Lifespan Machines, at which time monitoring of survivorship began. Deaths were not scored manually in the assays prior to loading plates onto the Automated Lifespan Machines. Scanner data were collected and analyzed using the previously published Automated Lifespan Machine software (https://github.com/nstroustrup/lifespan; (Stroustrup *et al.* 2013)). To account for species specific differences, nine posture files were generated using the posture file generation function of the Lifespan Machine software. Posture files were specifically generated for three *C. elegans* strains (N2, MY16, JU775), three *C. briggsae* strains (AF16, JU1348, HK104), and three *C. tropicalis* strains (JU1370, QG834, JU1630). To generate the standardized posture files, all three CITP locations independently created files for each of the nine strains, and the posture files from the three locations for each strain were averaged to generate the strain specific posture file. For strains without a strain specific posture file, the species reference strain posture file was used (*C. elegans*=N2, *C. briggsae*=AF16, and *C. tropicalis*= JU1630).

At the completion of experiments, the images were analyzed using the ALM software using the strain specific posture files. To validate the machine calls we used the built in “storyboarding” function of the ALM software (see storyboarding SOP doi.org/10.6084/m9.figshare.c.4580546). Storyboarding allows the experimenter to review each potential worm death and exclude machine detected objects that are not worms or modify the machine call of time of death (Stroustrup *et al.* 2013).

### Chemical treatments

Chemical compound treatments were performed as previously described (Lucanic *et al.* 2017). The compounds used were resveratrol (Cayman Chemical Company 70675, lot # 0414330-182), thioflavin T (MP Biomedicals 156877, lot # M6490), propyl gallate (Sigma Aldrich P3130l, lot # MKBR8169V), NP1 (Chembridge Labs custom order), and α-ketoglutaric acid (Sigma Aldrich K1128, lot # BCBF0081V). All compounds were first obtained as solids and dissolved in either water or DMSO (dimethyl sulfoxide) to obtain stock solutions. Control plates for chemical treatments were always included and contained the solvent specific for that compound. Compounds were added to culture plates such that the final volume was presumed equal to the agar volume of the plate. DMSO was present at 0.25% in all assays in which it was included. The final concentration in the plates for the compounds were 8 mM AKG, 50 µM ThT, 200 µM Propyl gallate, 50 µM NP1, and 100 µM resveratrol. Treatments were performed by moving worms onto compound treated plates. All compounds were administered to worms chronically throughout the survival study, starting at the first day of adulthood. Plate transfers through the 5^th^ day of adulthood were the same for manual and ALM assays. For ALM assays, transfers ceased after the fifth day of adulthood when plates were were positioned on the scanner bed, while in manual assays transfers of adults to compound or control plates continued every 3-4 days.

### Scanner light exposure

The CITP SOP for automated lifespan analysis includes imaging one column of plates every 15 minutes, so that each of the four columns of plates are scanned hourly. Because the fluorescent bulb runs perpendicular to the column scanned, all four columns are illuminated during each scheduled image capture every 15 minutes. To address potential light effects due to the frequent light exposure, we developed a filtering protocol using commercially available tinted plastics (Roscolux Supergel R10 medium yellow). To filter the transmitted light on the scanners the filter body-colored polycarbonate sheets were cut to the size and attached to the glass on the EPSON V700 transparency unit.

The manufacturer specifications for the filters show that the filter blocks approximately 100% of passing light ranging from 400 through 460 nm. The scanners automatically adjust the scan duration to account for variable bulb intensities. This resulted in longer scan durations when the light was filtered, which interfered with the completion of a 15-minute scan schedule. We addressed the slow scan speed by increasing the inter-scan duration to 30 minutes, with each of the four columns of plates being scanned once every two hours. The changes in the lighting and scan schedule did not appear to compromise the capabilities of the hardware and ALM software to analyze images, track animals and calculate death times for survival.

### Statistical analyses

Statistical analyses were performed as previously published for the manual assay dataset (Lucanic *et al.* 2017). In brief, a mix-model approach was used in which compound treatment and assay type were treated as fixed effects, and the other potential factors were treated as random effects. To accomplish this, we analyzed longevity using both GLMs using the lme4 v.1.12 package and a mixed model Cox proportional hazard (CPH) model using the coxme v.2.2-5 package in the R statistical language.

To test for the effects of individual compounds, we used CPH analysis within each strain so that each compound treatment replicate could be matched with their appropriate replicate-specific control in the randomized blocks design. Compound effects were tested as a planned comparison between the responses of individuals raised on the compound in question and those raised on the appropriate carrier control (H_2_O or DMSO).

### Percent of observed deaths returned for each approach

After collecting our ALM dataset, we compared the return rates (deaths observed/total number of animals at experiment start) to assess and compare worm loss between ALM and manual experiments. With conventional lifespan assays, 35-40 worms are cultured on each plate, while the larger (50 vs 38 mm diameter) ALM plates had 45-55 worms cultured on each plate. Using the approximate values of 37.5 and 50 worms per plate, we estimated the mean number of death observations per plate for both assays (Online Resource 20). As expected, this value also varied dramatically across strains for both conventional and ALM assays. In conventional assays death observation percentages ranged from 61% of possible death observations made for NIC122 to 88% for JU1088. In ALM assays, the average return rate values were generally lower, ranging from just 40% in JU1630 to 76% in N2.

## Supporting information

Online Resource 1

Online Resource 2

Online Resource 3

Online Resource 4

Online Resource 5

Online Resource 6

Online Resource 7

Online Resource 8

Online Resource 9

Online Resource 10

Online Resource 11

Online Resource 12

Online Resource 13

Online Resource 14

Online Resource 15

Online Resource 16

Online Resource 17

Online Resource 18

Online Resource 19

Online Resource 20

## Acknowledgements

We acknowledge all of the members of the Lithgow, Driscoll and Phillips labs for helpful discussions. We thank the CITP Advisory Committee and Ronald Kohanski (National Institute on Aging) for extensive discussion. We thank Asher Cutter, Marie-Anne Félix, and Christian Braendle for providing strains that they had directly collected. We thank Allie Kirsch for assistance storyboarding. Finally, we thank Nick Stroustrup for helpful discussions on the ALMs. Additional strains were provided by the CGC, which is funded by NIH Office of Research Infrastructure Programs (P40 OD010440). This work was supported by funding from the Larry L. Hillblom Foundation, the Glenn Foundation for Medical Research and National Institutes of Health grants (UL1 RR024917, supporting the Interdisciplinary Research Consortium on Geroscience, R01 AG029631, R21 AG048528, U01 AG045844, U01 AG045864, U01 AG045829, U24 AG056052).

## Abbreviations

CITP: *Caenorhabditis* Intervention Testing Program
ALMs: Automated Lifespan Machines
ThT: Thioflavin T
AKG: α-ketoglutarate
GL: general linear
CPH: Cox proportional hazard

## References

Al-Maqdi K. A., S. M. Hisaindee, M. A. Rauf, and S. S. Ashraf, 2017 Comparative Degradation of a Thiazole Pollutant by an Advanced Oxidation Process and an Enzymatic Approach. Biomolecules 7. https://doi.org/10.3390/biom7030064

Alavez S., M. C. Vantipalli, D. J. S. Zucker, I. M. Klang, and G. J. Lithgow, 2011 Amyloid-binding compounds maintain protein homeostasis during ageing and extend lifespan. Nature 472: 226–9. https://doi.org/10.1038/nature09873

Banse S. A., B. W. Blue, K. J. Robinson, C. M. Jarrett, and P. C. Phillips, 2019 The Stress-Chip: A microfluidic platform for stress analysis in Caenorhabditis elegans. PLoS One 14: e0216283. https://doi.org/10.1371/journal.pone.0216283

Castillo-Quan J. I., K. J. Kinghorn, and I. Bjedov, 2015 Genetics and pharmacology of longevity: the road to therapeutics for healthy aging. Adv. Genet. 90: 1–101. https://doi.org/10.1016/bs.adgen.2015.06.002

Chin R. M., X. Fu, M. Y. Pai, L. Vergnes, H. Hwang, et al., 2014 The metabolite α-ketoglutarate extends lifespan by inhibiting ATP synthase and TOR. Nature 510: 397–401. https://doi.org/10.1038/nature13264

Harrison D. E., R. Strong, Z. D. Sharp, J. F. Nelson, C. M. Astle, et al., 2009 Rapamycin fed late in life extends lifespan in genetically heterogeneous mice. Nature 460: 392–5. https://doi.org/10.1038/nature08221

Lithgow G. J., M. Driscoll, and P. Phillips, 2017 A long journey to reproducible results. Nature 548: 387–388. https://doi.org/10.1038/548387a

Lucanic M., T. Garrett, I. Yu, F. Calahorro, A. Asadi Shahmirzadi, et al., 2016 Chemical activation of a food deprivation signal extends lifespan. Aging Cell 15: 832–41. https://doi.org/10.1111/acel.12492

Lucanic M., W. T. Plummer, E. Chen, J. Harke, A. C. Foulger, et al., 2017 Impact of genetic background and experimental reproducibility on identifying chemical compounds with robust longevity effects. Nat. Commun. 8: 14256. https://doi.org/10.1038/ncomms14256

Magalhaes Filho C. D. De, B. Henriquez, N. E. Seah, R. M. Evans, L. R. Lapierre, et al., 2018 Visible light reduces *C. elegans* longevity. Nat. Commun. 9. https://doi.org/10.1038/s41467-018-02934-5

Maglioni S., N. Arsalan, and N. Ventura, 2016 *C. elegans* screening strategies to identify pro-longevity interventions. Mech. Ageing Dev. 157: 60–9. https://doi.org/10.1016/j.mad.2016.07.010

Maures T. J., L. N. Booth, B. A. Benayoun, Y. Izrayelit, F. C. Schroeder, et al., 2014 Males shorten the life span of *C. elegans* hermaphrodites via secreted compounds. Science 343: 541–4. https://doi.org/10.1126/science.1244160

Mishur R. J., M. Khan, E. Munkácsy, L. Sharma, A. Bokov, et al., 2016 Mitochondrial metabolites extend lifespan. Aging Cell 15: 336–48. https://doi.org/10.1111/acel.12439

Onken B., and M. Driscoll, 2010 Metformin induces a dietary restriction-like state and the oxidative stress response to extend *C. elegans* Healthspan via AMPK, LKB1, and SKN-1. PLoS One 5: e8758. https://doi.org/10.1371/journal.pone.0008758

Petrascheck M., and D. L. Miller, 2017 Computational Analysis of Lifespan Experiment Reproducibility. Front. Genet. 8: 92. https://doi.org/10.3389/fgene.2017.00092

Plummer W. T., J. Harke, M. Lucanic, E. Chen, D. Bhaumik, et al., 2017 Standardized Protocols from the *Caenorhabditis* Intervention Testing Program 2013-2016: Conditions and Assays used for Quantifying the Development, Fertility and Lifespan of Hermaphroditic *Caenorhabditis* Strains. Protoc. Exch. https://doi.org/10.1038/protex.2016.086

Shi C., and C. T. Murphy, 2014 Mating induces shrinking and death in *Caenorhabditis* mothers. Science 343: 536–40. https://doi.org/10.1126/science.1242958

Stroustrup N., B. E. Ulmschneider, Z. M. Nash, I. F. López-Moyado, J. Apfeld, et al., 2013 The *Caenorhabditis elegans* Lifespan Machine. Nat. Methods 10: 665–70. https://doi.org/10.1038/nmeth.2475

Stroustrup N., W. E. Anthony, Z. M. Nash, V. Gowda, A. Gomez, et al., 2016 The temporal scaling of *Caenorhabditis elegans* ageing. Nature 530: 103–7. https://doi.org/10.1038/nature16550

Wood J. G., B. Rogina, S. Lavu, K. Howitz, S. L. Helfand, et al., 2004 Sirtuin activators mimic caloric restriction and delay ageing in metazoans. Nature 430: 686–9. https://doi.org/10.1038/nature02789

Yoshimura J., K. Ichikawa, M. J. Shoura, K. L. Artiles, I. Gabdank, et al., 2019 Recompleting the *Caenorhabditis elegans* genome. Genome Res. 29: 1009–1022. https://doi.org/10.1101/gr.244830.118

