## Supplementary material for "Automated Lifespan Determination Across *Caenorhabditis* Strains and Species Reveals Assay-Specific Effects of Chemical Interventions": Online Resource 1

**Online Resource 1** Summary of ALM lifespan data under baseline conditions, and comparison to median lifespan from comparable manual assays

| | | ALM | | | | | | | | Manual | |
|---|---|---|---|---|---|---|---|---|---|---|---|
| **Species** | **Strain** | **Number of deaths** | **Number censored** | **Total observed** | **Mean lifespan** | **SEM** | **Median lifespan** | **Lower 95% CI** | **Upper 95% CI** | **Median lifespan** | **% diff median LS from manual** |
| ***C.elegans*** | N2 | 753 | 0 | 753 | 16.5 | 0.10 | 17.0 | 16.6 | 17.1 | 19 | -11 |
| | N2 PD1073 | 856 | 2 | 858 | 15.8 | 0.12 | 15.5 | 15.1 | 15.8 | - | - |
| | CB4856 | 628 | 0 | 628 | 17.9 | 0.16 | 18.2 | 17.6 | 18.5 | 20 | -9 |
| | ED3040 | 698 | 3 | 701 | 15.2 | 0.12 | 14.8 | 14.5 | 15.0 | 20 | -26 |
| | JU1088 | 848 | 1 | 849 | 17.0 | 0.13 | 17.0 | 16.7 | 17.4 | 18 | -5 |
| | JU1652 | 670 | 3 | 673 | 17.8 | 0.18 | 17.8 | 17.5 | 18.4 | 21 | -15 |
| | JU775 | 675 | 3 | 678 | 17.2 | 0.17 | 16.6 | 16.1 | 17.1 | 21 | -21 |
| | MY16 | 660 | 5 | 665 | 16.5 | 0.14 | 16.8 | 16.3 | 17.3 | 18 | -7 |
| | QX1211 | 180 | 0 | 180 | 16.6 | 0.26 | 15.9 | 15.5 | 16.7 | 19 | -16 |
| ***C. briggsae*** | AF16 | 398 | 6 | 404 | 20.0 | 0.24 | 20.0 | 19.4 | 20.7 | 25 | -20 |
| | ED3092 | 575 | 1 | 576 | 22.5 | 0.20 | 23.1 | 22.8 | 23.4 | 25 | -8 |
| | HK104 | 578 | 4 | 582 | 27.9 | 0.26 | 29.1 | 28.7 | 29.6 | 33 | -12 |
| | JU1264 | 580 | 0 | 580 | 21.2 | 0.28 | 20.3 | 19.5 | 21.2 | 23 | -12 |
| | JU1348 | 524 | 0 | 524 | 19.2 | 0.25 | 18.5 | 17.9 | 19.0 | 23 | -20 |
| | JU726 | 377 | 0 | 377 | 18.2 | 0.24 | 18.0 | 17.5 | 18.5 | 19 | -5 |
| | NIC20 | 637 | 2 | 639 | 18.4 | 0.17 | 18.3 | 17.8 | 18.8 | 23 | -20 |
| | QR25 | 758 | 3 | 761 | 18.3 | 0.19 | 17.2 | 16.9 | 17.7 | 18 | -4 |
| ***C. tropicalis*** | JU1373 | 566 | 3 | 569 | 18.0 | 0.10 | 18.3 | 18.1 | 18.5 | 23 | -21 |
| | JU1630 | 334 | 1 | 335 | 19.8 | 0.19 | 20.0 | 19.6 | 20.5 | 26 | -23 |
| | NIC122 | 268 | 0 | 268 | 15.9 | 0.28 | 16.8 | 15.5 | 17.5 | 22 | -23 |
| | NIC58 | 544 | 0 | 544 | 15.7 | 0.21 | 15.3 | 14.6 | 16.3 | 21 | -27 |
| | QG131 | 572 | 0 | 572 | 18.5 | 0.17 | 18.6 | 18.3 | 19.0 | 22 | -15 |
| | QG834 | 779 | 3 | 782 | 18.8 | 0.12 | 19.0 | 18.7 | 19.1 | 23 | -18 |
| | | | Combined total | 13,498 | | | | | | | |
