## Supplementary material for "Automated Lifespan Determination Across *Caenorhabditis* Strains and Species Reveals Assay-Specific Effects of Chemical Interventions": Online Resource 2

**Online Resource 2 ALM lifespan analysis results in shorter lived *C. elegans*, *C. briggsae* and *C. tropicalis* strains when compared to manual analysis.**

Survivorship curves for *C. elegans*, *C. briggsae* and *C. tropicalis* strains measured using manual (dashed line) or automated (solid line) analysis. ALM analysis consistently yields left-shifted lifespan curves. All comparisons are significantly different except *C. briggsae* strains JU726 ($p$=0.162), JU1264 ($p$=0.187), and QR25 ($p$=0.702). Comparisons were made using the CPH model.

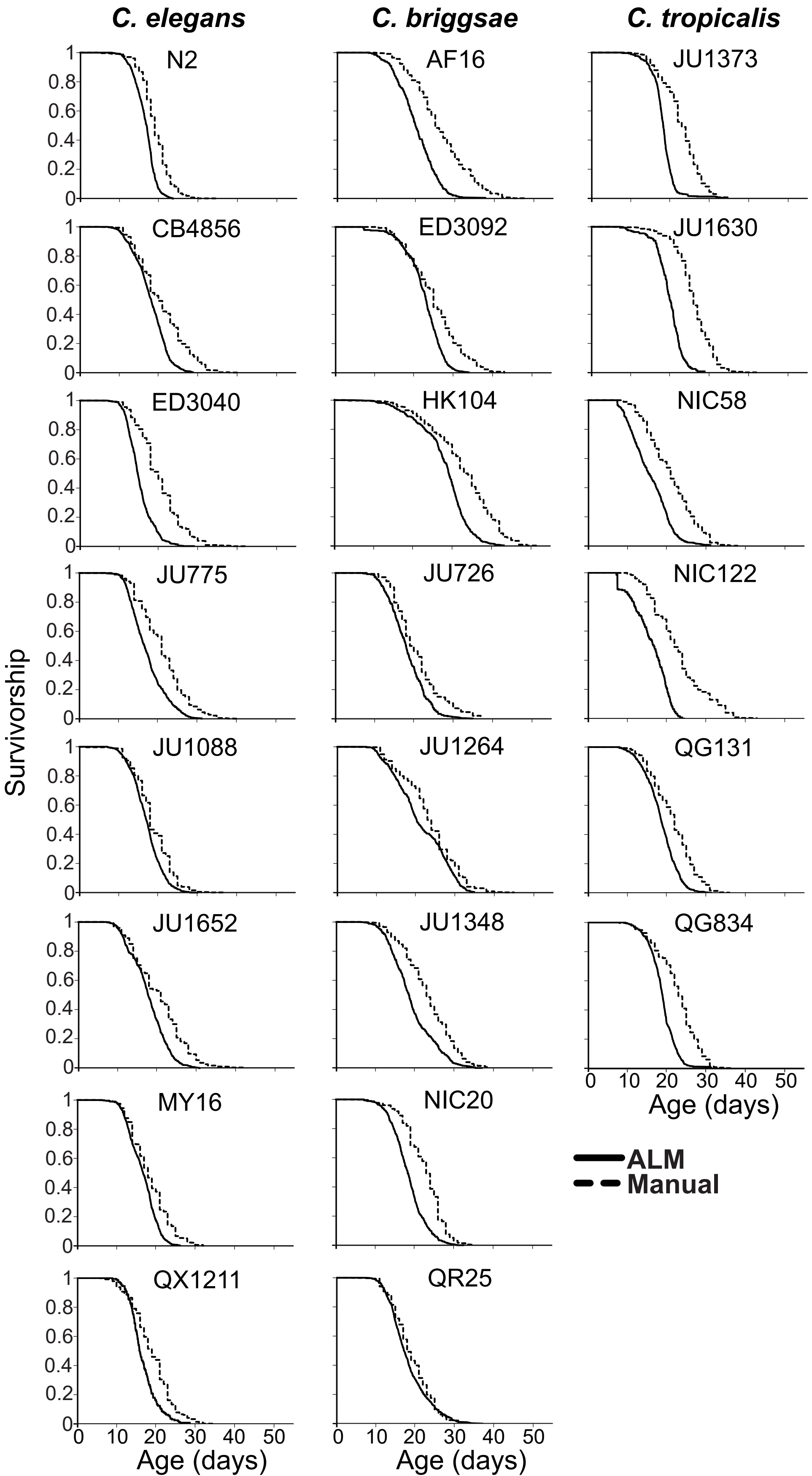

C. elegans
C. briggsae
C. tropicalis
N2
AF16
JU1373
CB4856
ED3092
JU1630
ED3040
HK104
NIC58
JU775
JU726
NIC122
JU1088
JU1264
QG131
JU1652
JU1348
QG834
MY16
NIC20
QX1211
QR25
Survivorship
Age (days)
ALM
Manual
