## Supplementary material for "Automated Lifespan Determination Across *Caenorhabditis* Strains and Species Reveals Assay-Specific Effects of Chemical Interventions": Online Resource 3

**Online Resource 3 Supplemental Text**

In survivorship assays, most non-death events result in an animal being censored from the study at the time at which the event is detected. These censored events are then included in the data set to calculate the survivor function and influence estimations of median lifespan and significance tests. These censored events are therefore important factors in survivorship studies and while the numbers vary, they can represent a large percent of animal observations in a data set. Analysis of the number of censored animals from our previous study (Lucanic *et al.* 2017) (Online Resource 5) indicated that 20% of the total animal observations were from censored animals. The percent of animals that were censored varied dramatically amongst strains and may be representative of their propensity to crawl off the plate, burrow under the agar, or experience a failure of egg-laying. Specifically, we observed a range from 8% censored in JU1652 to 36% censored in JU1264 (Online Resource 5). In our use of the Lifespan Machine (which we abbreviate ALM, for Automated Lifespan Machine), we do not record censored animals. Only animals observed to have died are recorded in our ALM data set. Therefore, unless otherwise noted, all comparisons between conventional and ALM datasets exclude censored animals.

*Percent of observed deaths returned for each approach*

After collecting our ALM dataset, we first compared the return rates (deaths observed) to assess whether more worms are lost from ALM or conventional experiments. With conventional lifespan assays, 35-40 worms are cultured on each plate, while the larger plates for the ALM had 45-55. Using the approximate values of 37.5 and 50 worms per plate, we estimated the amount of death observations per plate for both assays (Online Resource 20). As expected, this value also varied dramatically across strains for both conventional and ALM assays. In conventional assays this ranged from 61% of possible death observations made for NIC122 to 88% for JU1088. In ALM assays the average return rate values were generally lower, ranging from just 40% in JU1630 to 76% in N2.

**References**

Lucanic M., W. T. Plummer, E. Chen, J. Harke, A. C. Foulger, *et al.*, 2017 Impact of genetic background and experimental reproducibility on identifying chemical compounds with robust longevity effects. Nat. Commun. 8: 14256. https://doi.org/10.1038/ncomms14256
