## Supplementary material for "Automated Lifespan Determination Across *Caenorhabditis* Strains and Species Reveals Assay-Specific Effects of Chemical Interventions": Online Resource 4

**Online Resource 4 Survival differences among *C. elegans*, *C. briggsae* and *C. tropicalis* species are not due to differences in censoring**

(A) Survivorship curves for combined manual trials of each species indicated, when animals that died of unnatural causes were censored or (B) when all animals, including the censored population, were included in data analysis. (C) The basic shape and relative order of survival are maintained among species when data from all strains generated on the automated lifespan machines are combined. We did observe some strain-specific differences within species (see Online Resource 2).

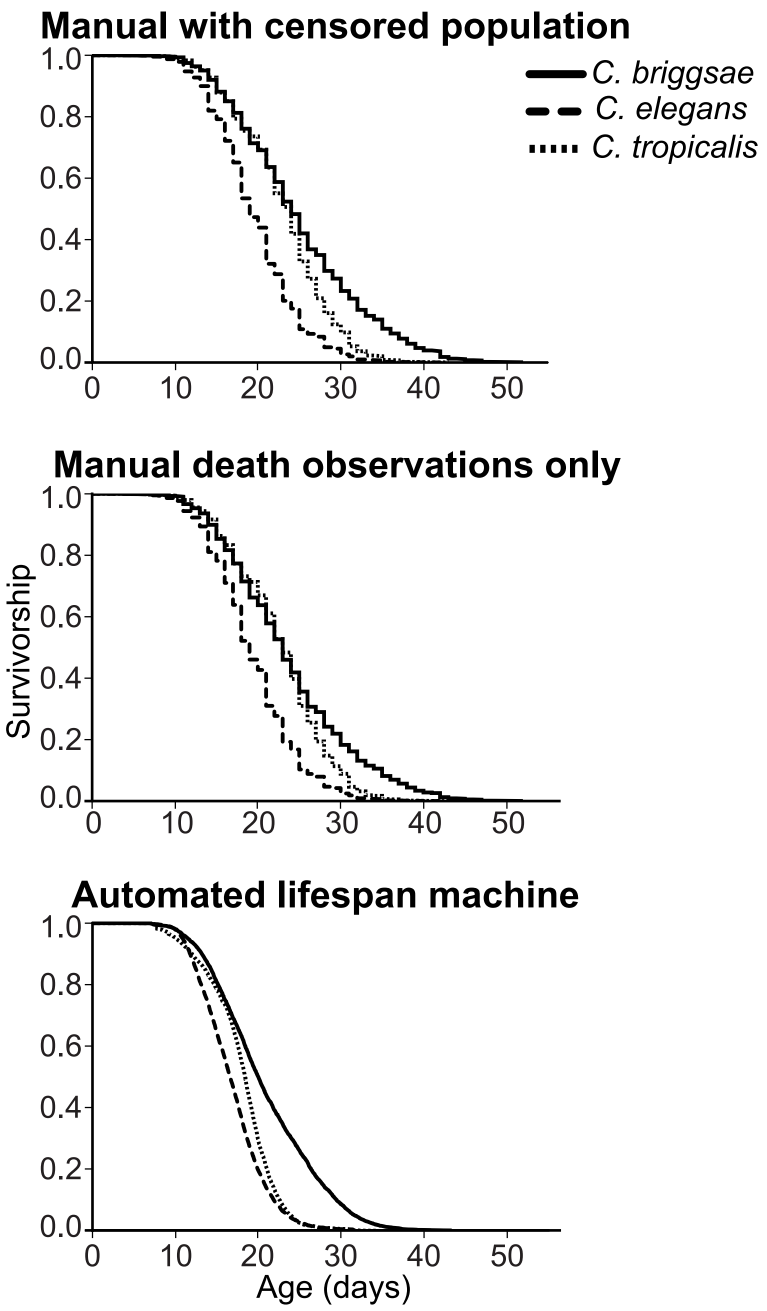

Manual with censored population
C. briggsae
C. elegans
C. tropicalis
Manual death observations only
Automated lifespan machine
Survivorship
Age (days)
1.0
0.8
0.6
0.4
0.2
0.0
0
10
20
30
40
50
