## Supplementary material for "Automated Lifespan Determination Across *Caenorhabditis* Strains and Species Reveals Assay-Specific Effects of Chemical Interventions": Online Resource 5

**Online Resource 5** Summary of manual dataset: deaths and censored observations. Strains are ordered from the smallest to largest percent censored.

| Species | Strain | Observed deaths | Number censored | Total observations | Percent censored |
|---|---|---|---|---|---|
| *C. elegans* | JU1652 | 1070 | 98 | 1168 | 8 |
| *C. elegans* | JU188 | 1081 | 107 | 1188 | 9 |
| *C. elegans* | ED3040 | 1047 | 111 | 1158 | 10 |
| *C. elegans* | JU775 | 1220 | 153 | 1373 | 11 |
| *C. elegans* | N2 | 3026 | 404 | 3430 | 12 |
| *C. elegans* | QX1211 | 1099 | 147 | 1246 | 12 |
| *C. elegans* | CB4856 | 1034 | 141 | 1175 | 12 |
| *C. tropicalis* | JU1630 | 826 | 124 | 950 | 13 |
| *C. elegans* | MY16 | 1112 | 175 | 1287 | 14 |
| *C. tropicalis* | JU1373 | 908 | 145 | 1053 | 14 |
| *C. tropicalis* | QG834 | 716 | 137 | 853 | 16 |
| *C. briggsae* | JU726 | 783 | 185 | 968 | 19 |
| *C. tropicalis* | QG131 | 683 | 227 | 910 | 25 |
| *C. briggsae* | NIC20 | 755 | 305 | 1060 | 29 |
| *C. tropicalis* | NIC58 | 603 | 246 | 849 | 29 |
| *C. tropicalis* | NIC122 | 620 | 272 | 892 | 30 |
| *C. briggsae* | AF16 | 904 | 398 | 1302 | 31 |
| *C. briggsae* | HK104 | 954 | 436 | 1390 | 31 |
| *C. briggsae* | QR25 | 670 | 316 | 986 | 32 |
| *C. briggsae* | ED3092 | 666 | 325 | 991 | 33 |
| *C. briggsae* | JU1348 | 649 | 331 | 980 | 34 |
| *C. briggsae* | JU1264 | 717 | 407 | 1124 | 36 |
| | Combined | 21143 | 5190 | 26333 | 20 |
