## Supplementary material for "Automated Lifespan Determination Across *Caenorhabditis* Strains and Species Reveals Assay-Specific Effects of Chemical Interventions": Online Resource 6

**Online Resource 6** Partitioning of variation for longevity across genetic background and experimental replicates within and among labs. Variance components were estimated as a randomized block design using a restricted maximum likelihood (REML) general linear model using the *lme4* package of R. All factors were treated as random effects. Tests involving the fixed compound effects are reported in Online Resource 9.

A. Variance component estimates for the natural log of individual lifespan from the 22-strain, three-species baseline longevity experiments. Sample sizes and summary statistics are presented in Online Resource 1.

| Source | Variance Component |
|---|---|
| Species | 1.66E-03 |
| Strain*Species | 1.80E-03 |
| Lab | 3.08E-04 |
| Lab*Species | 9.26E-05 |
| Lab*Strain | 1.10E-03 |
| Scanner*Lab | 6.01E-09 |
| Trial[Scanner, Lab] | 3.98E-04 |
| Plate_Trial[Trial, Scanner, Lab] | 1.28E-03 |
| Residual | 9.29E-03 |
| Total | 1.59E-02 |

B. Average variance component estimates for the natural log of individual lifespan from the six-strain, three-compound experiments. See Online Resource 14 for a per-strain analysis of replication and samples sizes. Summary statistics are presented in Online Resource 11.

| Source | Variance Component |
|---|---|
| Species | 2.42E-03 |
| Strain*Species | 2.51E-03 |
| Compound*Species | 9.58E-04 |
| Compound*Strain | 3.75E-04 |
| Lab | 0.00E+00 |
| Lab*Species | 0.00E+00 |
| Lab*Strain | 1.55E-03 |
| Compound*Lab | 2.32E-05 |
| Scanner*Lab | 3.96E-10 |
| Trial[Scanner, Lab] | 5.34E-04 |
| Plate_Trial[Trial, Scanner, Lab] | 1.04E-03 |
| Residual | 9.52E-03 |
| Total | 1.89E-02 |

C. Average variance component estimates for the natural log of individual lifespan from the corresponding manual experiments. Sample sizes and summary statistics are presented in Online Resources 1 and 5.

| Source | Variance Component | Lower 95% CI | Upper 95% CI | Percent of Total |
|---|---|---|---|---|
| Species | 5.23E-02 | 0.00E+00 | 4.91E-01 | 32.5 |
| Strain*Species | 1.15E-02 | 3.08E-03 | 7.30E-02 | 7.1 |
| Compound*Species | 5.21E-03 | 1.79E-04 | 9.46E-03 | 3.2 |
| Compound*Strain | 3.26E-03 | 1.46E-03 | 7.78E-03 | 2.0 |
| Lab | 0.00E+00 | 0.00E+00 | 7.29E-03 | 0.0 |
| Lab*Species | 6.79E-04 | 0.00E+00 | 6.27E-03 | 0.4 |
| Lab*Strain | 3.94E-04 | 0.00E+00 | 2.06E-03 | 0.2 |
| Compound*Lab | 0.00E+00 | 0.00E+00 | 7.65E-04 | 0.0 |
| Experimenter*Lab | 2.34E-03 | 3.71E-04 | 7.37E-03 | 1.5 |
| Trial[Experimenter, Lab] | 3.46E-03 | 1.93E-03 | 6.06E-03 | 2.2 |
| Plate_Trial[Trial, Experimenter, Lab] | 8.29E-03 | 6.92E-03 | 9.90E-03 | 5.1 |
| Residual | 7.36E-02 | 7.19E-02 | 7.52E-02 | 45.7 |
| Total | 1.61E-01 | | | 100.0 |
