## Supplementary material for "Automated Lifespan Determination Across *Caenorhabditis* Strains and Species Reveals Assay-Specific Effects of Chemical Interventions": Online Resource 7

**Online Resource 7 Several lifespan compound interventions are reported similarly by manual and ALM analysis.**

The median lifespan under adult exposure to NP1 (a), propyl gallate (b) or resveratrol (c) are shown for three *C. elegans* (N2, JU775, and MY16) and *C. briggsae* (AF16, JU1348, and HK104) strains. Each point represents the median lifespan from an individual plate trial. The bars represent the mean +/- the standard error of the mean. Replicates were generated at the three CITP sites (Blue-Buck Institute, Green-Oregon and Red- Rutgers). Lifespans were measured for compound (circles) and vehicle control (triangles) conditions. Asterisks represent $p$-values (****$p$<0.0001, *** $p$<0.001, ** $p$<0.01 and * $p$<0.05) from the CPH model when comparing the lifespans under compound exposure versus the lifespans exposed to the vehicle control.

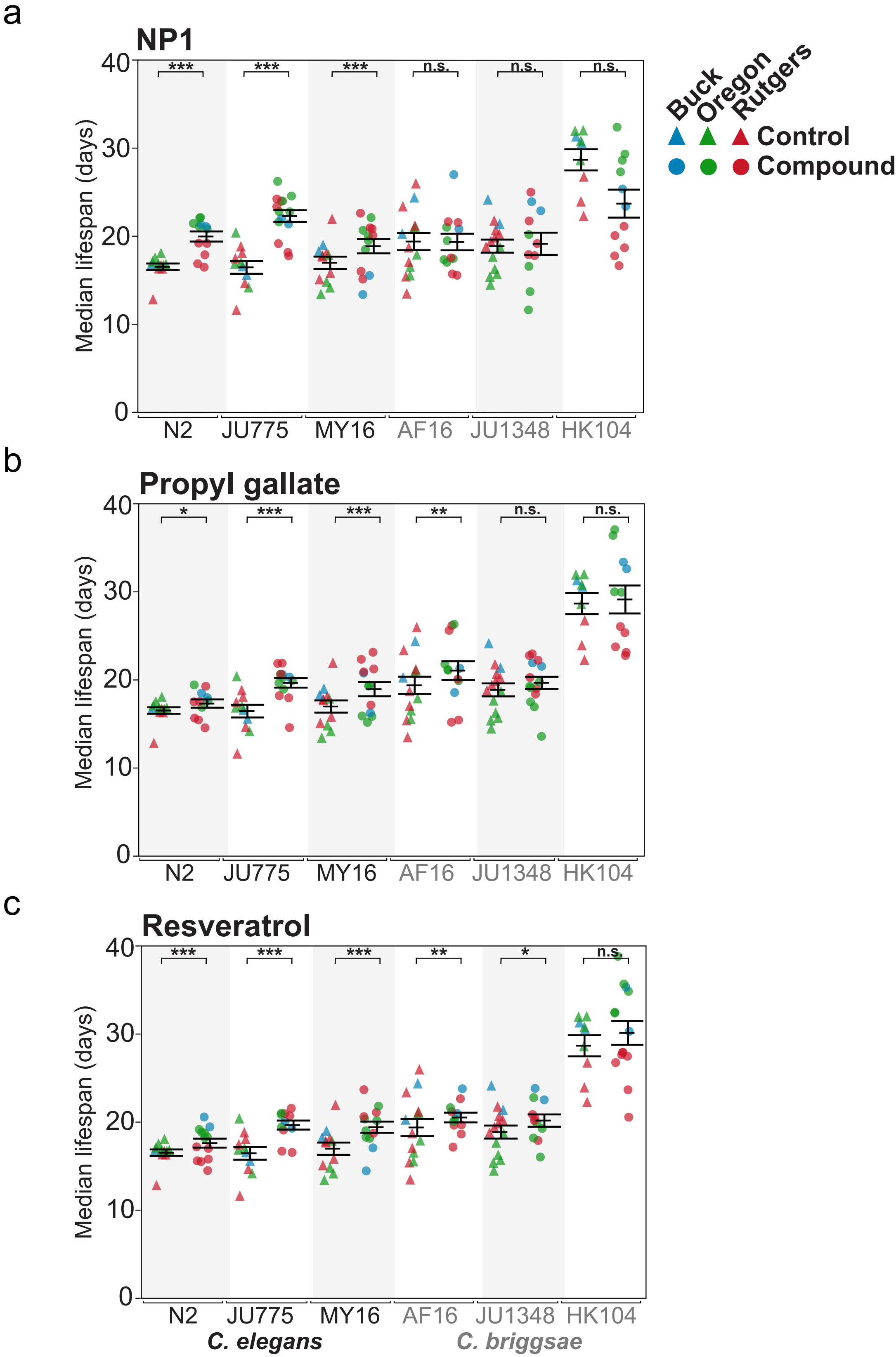

a
NP1
b
Propyl gallate
c
Resveratrol
Median lifespan (days)
0
10
20
30
40
***
n.s.
**
*
N2
JU775
MY16
AF16
JU1348
HK104
C. elegans
C. briggsae
Buck
Oregon
Rutgers
Control
Compound
