## Supplementary material for "Automated Lifespan Determination Across *Caenorhabditis* Strains and Species Reveals Assay-Specific Effects of Chemical Interventions": Online Resource 8

**Online Resource 8 Thioflavin T, but not α-ketoglutarate, lifespan effects are reversed by light exposure in ALM analysis.**

The median lifespans under adult exposure to thioflavin T (a) or α-ketoglutarate (b) are shown for three *C. elegans* (N2, JU775, and MY16) and *C. briggsae* (AF16, JU1348, and HK104) strains. Each point represents the median lifespan from an individual plate trial. The bars represent the mean +/- the standard error of the mean. Replicates were generated at three CITP sites (Blue-Buck Institute, Green-Oregon and Red- Rutgers). Lifespans were measured by standard automated survival analysis (vehicle control-triangles or compound - circles) or with automated lifespan analysis modified to accommodate light filtering (see materials and methods) (vehicle control-inverted triangles or compound-diamonds). Asterisks represent *p*-values (****$p<0.0001$, *** $p<0.001$, ** $p<0.01$ and * $p<0.05$) from the CPH model for the shown comparisons.

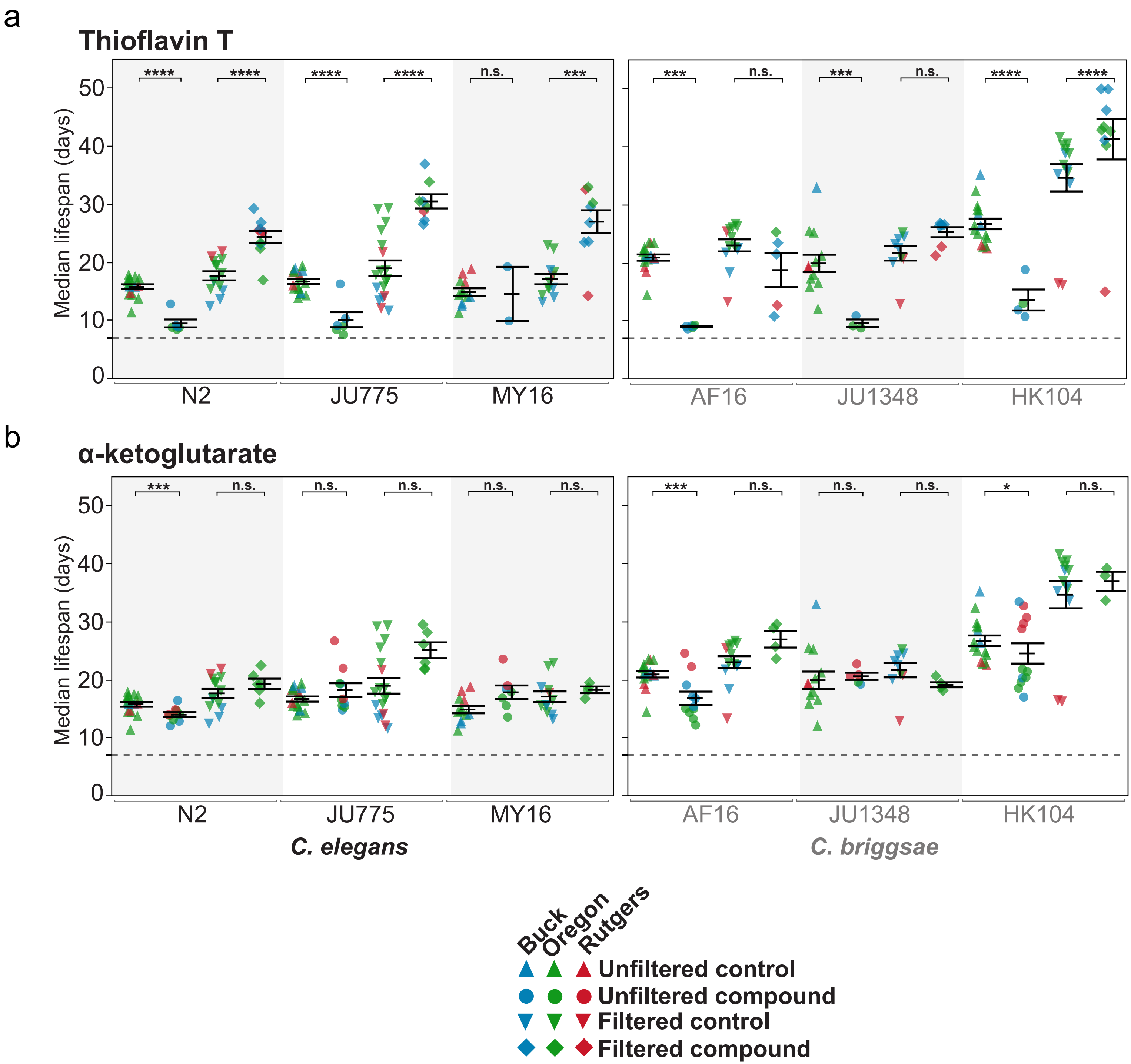
a
Thioflavin T
Median lifespan (days)
0
10
20
30
40
50
****
****
****
****
n.s.
***
***
n.s.
***
n.s.
****
****
N2
JU775
MY16
AF16
JU1348
HK104
b
α-ketoglutarate
Median lifespan (days)
***
n.s.
n.s.
n.s.
n.s.
n.s.
***
n.s.
n.s.
n.s.
*
n.s.
N2
JU775
MY16
AF16
JU1348
HK104
C. elegans
C. briggsae
Buck
Oregon
Rutgers
Unfiltered control
Unfiltered compound
Filtered control
Filtered compound
