## Supplementary material for "Automated Lifespan Determination Across *Caenorhabditis* Strains and Species Reveals Assay-Specific Effects of Chemical Interventions": Online Resource 9

**Online Resource 9** Manual vs. ALM: approximate average return rates per plate (observed mean number of deaths versus expected number of animals at experiment start) for compound trials. The expected number of animals per plate for manual assays was 37.5, and for ALM assays was 50.

| | | | Manual | | ALM | |
|---|---|---|---|---|---|---|
| **Species** | **Strain** | **Compound** | **Average obs. deaths** | **% expected observed** | **Average obs. deaths** | **% expected observed** |
| ***C. elegans*** | N2 | CTRL-H2O | 33 | 88 | 32 | 64 |
| | | CTRL-DMSO | 31 | 83 | 37 | 74 |
| | | AKG | 30 | 80 | 36 | 72 |
| | | NP1 | 32 | 85 | 39 | 78 |
| | | Propyl gallate | 29 | 77 | 37 | 74 |
| | | Resveratrol | 30 | 80 | 37 | 74 |
| | | ThT | 30 | 80 | 25 | 50 |
| | JU775 | CTRL-H2O | 32 | 85 | 26 | 52 |
| | | CTRL-DMSO | 29 | 77 | 35 | 70 |
| | | AKG | 25 | 67 | 30 | 60 |
| | | NP1 | 29 | 77 | 35 | 70 |
| | | Propyl gallate | 27 | 72 | 35 | 70 |
| | | Resveratrol | 28 | 75 | 33 | 66 |
| | | ThT | 27 | 72 | 19 | 38 |
| | MY16 | CTRL-H2O | 30 | 80 | 26 | 52 |
| | | CTRL-DMSO | 26 | 69 | 30 | 60 |
| | | AKG | 25 | 67 | 26 | 52 |
| | | NP1 | 27 | 72 | 33 | 66 |
| | | Propyl gallate | 24 | 64 | 29 | 58 |
| | | Resveratrol | 22 | 59 | 30 | 60 |
| | | ThT | 27 | 72 | 22 | 44 |
| ***C. briggsae*** | AF16 | CTRL-H2O | 20 | 53 | 19 | 38 |
| | | CTRL-DMSO | 21 | 56 | 21 | 42 |
| | | AKG | 17 | 45 | 15 | 30 |
| | | NP1 | 18 | 48 | 23 | 46 |
| | | Propyl gallate | 21 | 56 | 19 | 38 |
| | | Resveratrol | 17 | 45 | 20 | 40 |
| | | ThT | 19 | 51 | 20 | 40 |
| | JU1348 | CTRL-H2O | 19 | 51 | 21 | 42 |
| | | CTRL-DMSO | 20 | 53 | 23 | 46 |
| | | AKG | 20 | 53 | 21 | 42 |
| | | NP1 | 23 | 61 | 25 | 50 |
| | | Propyl gallate | 22 | 59 | 24 | 48 |
| | | Resveratrol | 19 | 51 | 28 | 56 |
| | | ThT | 18 | 48 | 24 | 48 |
| | HK104 | CTRL-H2O | 28 | 75 | 31 | 62 |
| | | CTRL-DMSO | 26 | 69 | 34 | 68 |
| | | AKG | 24 | 64 | 27 | 54 |
| | | NP1 | 28 | 75 | 36 | 72 |
| | | Propyl gallate | 25 | 67 | 33 | 66 |
| | | Resveratrol | 27 | 72 | 33 | 66 |
| | | ThT | 24 | 64 | 23 | 46 |
