## Supplementary material for "Automated Lifespan Determination Across *Caenorhabditis* Strains and Species Reveals Assay-Specific Effects of Chemical Interventions": Online Resource 10

**Online Resource 10** Summary of ALM lifespan data under compound treatment (NP1, propyl gallate, and resveratrol) conditions, and comparison to median lifespan from comparable manual assays

| | | | ALM | | | | | | | | Manual | |
|---|---|---|---|---|---|---|---|---|---|---|---|---|
| **Species** | **Strain** | **Compound** | **Number of deaths** | **Number censored** | **Total observed** | **Mean lifespan** | **SEM** | **Median lifespan** | **Lower 95% CI** | **Upper 95% CI** | **Median lifespan** | **% diff med LS from manual** |
| ***C. elegans*** | JU775 | CTRL-DMSO | 382 | 0 | 382 | 16.8 | 0.23 | 16.6 | 15.8 | 17.4 | 17 | -2 |
| | | NP1 | 492 | 0 | 492 | 21.6 | 0.20 | 22.3 | 21.7 | 22.7 | 21 | 6 |
| | | Propyl gallate | 459 | 0 | 459 | 19.5 | 0.22 | 20.0 | 19.3 | 20.6 | 23 | -13 |
| | | Resveratrol | 362 | 0 | 362 | 19.9 | 0.27 | 19.8 | 18.8 | 20.5 | 24 | -18 |
| | MY16 | CTRL-DMSO | 359 | 1 | 360 | 17.2 | 0.18 | 17.3 | 16.6 | 17.7 | 15 | 16 |
| | | NP1 | 441 | 0 | 441 | 18.6 | 0.18 | 19.4 | 18.9 | 19.8 | 21 | -8 |
| | | Propyl gallate | 351 | 2 | 353 | 19.1 | 0.24 | 19.4 | 18.7 | 20.4 | 18 | 8 |
| | | Resveratrol | 385 | 0 | 385 | 19.5 | 0.21 | 20.1 | 19.6 | 20.5 | 18 | 12 |
| | N2 | CTRL-DMSO | 444 | 0 | 444 | 16.4 | 0.12 | 16.8 | 16.6 | 17.0 | 17 | -1 |
| | | NP1 | 472 | 0 | 472 | 19.9 | 0.16 | 20.6 | 20.3 | 20.8 | 21 | -2 |
| | | Propyl gallate | 407 | 0 | 407 | 17.4 | 0.16 | 17.5 | 17.0 | 17.7 | 19 | -8 |
| | | Resveratrol | 483 | 0 | 483 | 17.6 | 0.15 | 17.7 | 17.3 | 18.1 | 19 | -7 |
| ***C. briggsae*** | AF16 | CTRL-DMSO | 299 | 0 | 299 | 19.4 | 0.30 | 18.7 | 18.1 | 19.6 | 26 | -28 |
| | | NP1 | 278 | 0 | 278 | 19.7 | 0.32 | 19.5 | 18.4 | 20.2 | 26 | -25 |
| | | Propyl gallate | 232 | 0 | 232 | 21.9 | 0.39 | 21.8 | 20.6 | 23.0 | 24 | -9 |
| | | Resveratrol | 219 | 1 | 220 | 20.7 | 0.38 | 20.7 | 19.6 | 21.5 | 24 | -14 |
| | HK104 | CTRL-DMSO | 303 | 1 | 304 | 28.9 | 0.32 | 30.1 | 29.4 | 30.6 | 24 | 25 |
| | | NP1 | 399 | 0 | 399 | 24.0 | 0.39 | 24.6 | 23.6 | 25.3 | 28 | -12 |
| | | Propyl gallate | 358 | 4 | 362 | 29.7 | 0.35 | 30.4 | 29.7 | 31.0 | 35 | -13 |
| | | Resveratrol | 466 | 4 | 470 | 30.0 | 0.35 | 31.0 | 30.3 | 31.6 | 35 | -11 |
| | JU1348 | CTRL-DMSO | 319 | 1 | 320 | 19.2 | 0.28 | 18.5 | 17.6 | 19.3 | 25 | -26 |
| | | NP1 | 278 | 0 | 278 | 19.7 | 0.35 | 19.4 | 18.9 | 20.2 | 28 | -31 |
| | | Propyl gallate | 333 | 0 | 333 | 20.1 | 0.30 | 19.7 | 19.2 | 20.3 | 25 | -21 |
| | | Resveratrol | 310 | 1 | 311 | 20.8 | 0.28 | 20.1 | 19.4 | 20.9 | 26 | -23 |
