## Supplementary material for "Automated Lifespan Determination Across *Caenorhabditis* Strains and Species Reveals Assay-Specific Effects of Chemical Interventions": Online Resource 11

**Online Resource 11** Significance tests for compound interventions effects on longevity. Each effect is tested using both a general linear model of age at death and random effects Cox Proportional Hazard Model. Each compound is tested as a planned comparison against its appropriate carrier control. Variance components estimates for the randomized-block effects that were included in the overall model are presented in Online Resources 14-17.

**A.** ***C. elegans*** **N2**

| | General linear model | | | | Random effects Cox Proportional Hazard | | | |
|---|---|---|---|---|---|---|---|---|
| Compound | Effect | Std err | *z*-value | p-value | Effect | Std err | *z*-value | *p*-value |
| NP1 | 3.74 | 0.33 | 11.39 | <1E-04 | -1.41 | 0.14 | -10.21 | <1E-04 |
| Propyl gallate | 1.32 | 0.36 | 3.65 | 0.0007 | -0.56 | 0.15 | -3.70 | 0.0006 |
| Resveratrol | 1.64 | 0.32 | 5.10 | <1E-04 | -0.77 | 0.13 | -5.78 | <1E-04 |
| Thio T, unfiltered | -5.96 | 0.96 | -6.23 | <1E-09 | 2.79 | 0.23 | 12.01 | <1E-10 |
| Thio T, filtered | 5.31 | 0.82 | 6.48 | <1E-10 | -1.18 | 0.19 | -6.36 | <1E-09 |
| AKG, unfiltered | -2.14 | 0.57 | -3.75 | <0.001 | 1.23 | 0.26 | 4.80 | <0.001 |
| AKG, pH adjusted | -0.68 | 0.46 | -1.50 | 0.2800 | 0.62 | 0.23 | 2.72 | 0.0163 |
| AKG, unfilt, Phillips | -2.11 | 0.78 | -2.69 | 0.0328 | 1.41 | 0.22 | 6.34 | <0.001 |
| AKG, filt, Phillips | 1.02 | 0.74 | 1.38 | 0.4865 | 0.14 | 0.21 | 0.67 | 0.9009 |
| AKG, pH adjusted, Phillips | -0.79 | 0.85 | -0.93 | 0.7730 | 0.35 | 0.24 | 1.49 | 0.4184 |

**B.** ***C. elegans*** **MY16**

| | General linear model | | | | Random effects Cox Proportional Hazard | | | |
|---|---|---|---|---|---|---|---|---|
| Compound | Effect | Std err | *z*-value | *p*-value | Effect | Std err | *z*-value | *p*-value |
| NP1 | 1.69 | 0.78 | 2.17 | 0.0754 | -0.45 | 0.24 | -1.85 | 0.1543 |
| Propyl gallate | 1.83 | 0.76 | 2.41 | 0.0408 | -0.86 | 0.24 | -3.59 | <0.001 |
| Resveratrol | 2.07 | 0.76 | 2.72 | 0.0169 | -0.77 | 0.24 | -3.26 | 0.0031 |
| Thio T, unfiltered | 1.03 | 1.86 | 0.56 | 0.8340 | -0.21 | 0.32 | -0.67 | 0.7665 |
| Thio T, filtered | 8.91 | 1.00 | 8.92 | <0.001 | -1.43 | 0.17 | -8.54 | <0.001 |
| AKG, unfiltered | 2.36 | 0.82 | 2.88 | 0.0102 | -0.33 | 0.24 | -1.40 | 0.3240 |
| AKG, pH adjusted | 0.19 | 0.64 | 0.30 | 0.9502 | 0.08 | 0.19 | 0.42 | 0.9050 |
| AKG, unfilt, Phillips | 1.04 | 0.70 | 1.49 | 0.4210 | -0.01 | 0.16 | -0.06 | 0.9999 |
| AKG, filt, Phillips | 0.27 | 0.54 | 0.51 | 0.9520 | 0.34 | 0.13 | 2.67 | 0.0348 |
| AKG, pH adjusted, Phillips | -0.52 | 0.53 | -0.98 | 0.7390 | 0.17 | 0.12 | 1.37 | 0.4944 |

**C. *C. elegans* JU775**

| | General linear model | | | | Random effects Cox Proportional Hazard | | | |
|---|---|---|---|---|---|---|---|---|
| Compound | Effect | Std err | *z*-value | *p*-value | Effect | Std err | *z*-value | *p*-value |
| NP1 | 4.85 | 0.50 | 9.70 | <1E-07 | -1.04 | 0.14 | -7.59 | <1E-05 |
| Propyl gallate | 2.84 | 0.51 | 5.58 | <1E-07 | -0.71 | 0.14 | -5.08 | <1E-05 |
| Resveratrol | 3.12 | 0.55 | 5.68 | <1E-07 | -0.94 | 0.15 | -6.22 | <1E-05 |
| Thio T, unfiltered | -6.63 | 1.53 | -4.32 | 0.0000 | 2.55 | 0.32 | 7.92 | <1E-06 |
| Thio T, filtered | 10.16 | 1.23 | 8.24 | <1E-05 | -1.20 | 0.26 | -4.68 | <1E-05 |
| AKG, unfiltered | 1.05 | 0.95 | 1.11 | 0.4950 | -0.01 | 0.22 | -0.04 | 0.9990 |
| AKG, pH adjusted | -1.34 | 0.83 | -1.62 | 0.2250 | 0.34 | 0.19 | 1.85 | 0.1450 |
| AKG, unfilt, Phillips | 0.08 | 1.13 | 0.07 | 0.9999 | 0.15 | 0.22 | 0.68 | 0.8951 |
| AKG, filt, Phillips | 1.82 | 0.94 | 1.94 | 0.1951 | 0.00 | 0.19 | 0.01 | 1.0000 |
| AKG, pH adjusted, Phillips | -3.46 | 0.96 | -3.61 | 0.0021 | 0.69 | 0.22 | 3.21 | 0.0064 |

**D. *C. briggsae* AF16**

| | General linear model | | | | Random effects Cox Proportional Hazard | | | |
|---|---|---|---|---|---|---|---|---|
| Compound | Effect | Std err | *z*-value | *p*-value | Effect | Std err | *z*-value | *p*-value |
| NP1 | 0.49 | 0.88 | 0.56 | 0.9066 | -0.15 | 0.16 | -0.94 | 0.6828 |
| Propyl gallate | 1.94 | 0.88 | 2.21 | 0.0733 | -0.48 | 0.16 | -2.94 | 0.0093 |
| Resveratrol | 1.75 | 0.90 | 1.95 | 0.1324 | -0.52 | 0.17 | -3.08 | 0.0061 |
| Thio T, unfiltered | -12.09 | 1.04 | -11.62 | <1E-04 | 5.33 | 0.34 | 15.72 | <1E-05 |
| Thio T, filtered | -1.04 | 1.22 | -0.85 | 0.6590 | -0.15 | 0.27 | -0.58 | 0.8210 |
| AKG, unfiltered | -3.79 | 0.78 | -4.88 | <1E-04 | 1.12 | 0.18 | 6.11 | <1E-04 |
| AKG, pH adjusted | -3.47 | 0.74 | -4.70 | <1E-04 | 0.96 | 0.17 | 5.65 | <1E-04 |
| AKG, unfilt, Phillips | -6.04 | 0.84 | -7.15 | <0.001 | 1.65 | 0.19 | 8.50 | <0.001 |
| AKG, filt, Phillips | 1.78 | 0.80 | 2.23 | 0.1055 | -0.29 | 0.18 | -1.65 | 0.3276 |
| AKG, pH adjusted, Phillips | -3.11 | 0.57 | -5.43 | <0.001 | 0.94 | 0.13 | 7.33 | <0.001 |

**E. *C. briggsae* HK104**

| | General linear model | | | | Random effects Cox Proportional Hazard | | | |
|---|---|---|---|---|---|---|---|---|
| Compound | Effect | Std err | *z*-value | *p*-value | Effect | Std err | *z*-value | *p*-value |
| NP1 | -4.04 | 0.90 | -4.48 | <0.001 | 0.52 | 0.20 | 2.58 | 0.0256 |
| Propyl gallate | 1.67 | 0.89 | 1.89 | 0.1404 | -0.35 | 0.19 | -1.80 | 0.1684 |
| Resveratrol | 2.10 | 0.86 | 2.44 | 0.0376 | -0.56 | 0.19 | -2.97 | 0.0081 |
| Thio T, unfiltered | -14.71 | 2.12 | -6.96 | <0.001 | 3.64 | 0.40 | 9.15 | <0.001 |
| Thio T, filtered | 3.81 | 1.46 | 2.61 | 0.0227 | -0.65 | 0.26 | -2.53 | 0.0283 |
| AKG, unfiltered | -3.29 | 1.35 | -2.44 | 0.0367 | 0.98 | 0.32 | 3.04 | 0.0061 |
| AKG, pH adjusted | -7.16 | 1.26 | -5.68 | <0.001 | 1.62 | 0.31 | 5.21 | <0.001 |
| AKG, unfilt, Phillips | -5.85 | 1.26 | -4.64 | <0.001 | 1.67 | 0.29 | 5.68 | <0.001 |
| AKG, filt, Phillips | -4.01 | 1.64 | -2.44 | 0.0633 | 0.51 | 0.37 | 1.37 | 0.4940 |
| AKG, pH adjusted, Phillips | -5.76 | 1.26 | -4.57 | <0.001 | 1.10 | 0.29 | 3.78 | <0.001 |

**F. *C. briggsae* JU1348**

| | General linear model | | | | Random effects Cox Proportional Hazard | | | |
|---|---|---|---|---|---|---|---|---|
| Compound | Effect | Std err | *z*-value | *p*-value | Effect | Std err | *z*-value | *p*-value |
| NP1 | 0.32 | 0.64 | 0.51 | 0.9274 | -0.20 | 0.14 | -1.39 | 0.3757 |
| Propyl gallate | 0.90 | 0.61 | 1.48 | 0.3234 | -0.24 | 0.14 | -1.73 | 0.0118 |
| Resveratrol | 1.60 | 0.62 | 2.60 | 0.0263 | -0.39 | 0.14 | -2.86 | 0.2064 |
| Thio T, unfiltered | -11.98 | 1.61 | -7.44 | <0.001 | 6.16 | 0.35 | 17.65 | <0.001 |
| Thio T, filtered | 2.61 | 1.19 | 2.20 | 0.0659 | -0.35 | 0.13 | -2.72 | 0.0155 |
| AKG, unfiltered | 0.69 | 1.59 | 0.43 | 0.8979 | -0.26 | 0.34 | -0.78 | 0.7006 |
| AKG, pH adjusted | -4.52 | 1.17 | -3.86 | <0.001 | 1.10 | 0.26 | 4.24 | <0.001 |
| AKG, unfilt, Phillips | 0.68 | 2.76 | 0.25 | 0.9940 | -0.16 | 0.55 | -0.29 | 0.9909 |
| AKG, filt, Phillips | -6.94 | 2.21 | -3.14 | 0.0082 | 1.28 | 0.45 | 2.83 | 0.0214 |
| AKG, pH adjusted, Phillips | -5.77 | 1.57 | -3.68 | 0.0011 | 1.25 | 0.32 | 3.94 | <0.001 |
