## Supplementary material for "Automated Lifespan Determination Across *Caenorhabditis* Strains and Species Reveals Assay-Specific Effects of Chemical Interventions": Online Resource 12

**Online Resource 12** Significance tests for compound interventions effects on longevity. Each effect is tested using both a general linear model of age at death and random effects Cox Proportional Hazard Model. Each compound is tested as a planned comparison against its appropriate carrier control. Variance components estimates for the randomized-block effects that were included in the overall model are presented in Online Resources 5-17.

**A.** ***C. elegans*** **N2**

| | General linear model | | | | Random effects Cox Proportional Hazard | | | |
|---|---|---|---|---|---|---|---|---|
| Compound | Effect | Std err | *z*-value | *p*-value | Effect | Std err | *z*-value | *p*-value |
| Thio T, filtered & unfiltered | -11.27 | 1.25 | -8.98 | <1E-10 | 3.97 | 0.30 | 13.33 | <1E-10 |
| AKG, unfiltered & pH adjusted | -1.45 | 0.73 | -1.99 | 0.1070 | 0.61 | 0.36 | 1.67 | 0.2007 |
| AKG, unfiltered & filtered | -3.13 | 1.08 | -2.90 | 0.0180 | 1.27 | 0.30 | 4.19 | <0.001 |
| AKG, pH adjusted & filtered | -1.81 | 1.13 | -1.60 | 0.3530 | 0.21 | 0.31 | 0.68 | 0.8950 |
| AKG, pH adjusted & unfiltered | 1.32 | 1.16 | 1.14 | 0.6449 | -1.06 | 0.32 | -3.28 | 0.0053 |

**B.** ***C. elegans*** **MY16**

| | General linear model | | | | Random effects Cox Proportional Hazard | | | |
|---|---|---|---|---|---|---|---|---|
| Compound | Effect | Std err | *z*-value | *p*-value | Effect | Std err | *z*-value | *p*-value |
| Thio T, filtered & unfiltered | -7.88 | 2.11 | -3.74 | <0.001 | 1.22 | 0.36 | 3.39 | 0.0017 |
| AKG, unfiltered & pH adjusted | 2.17 | 1.04 | 2.09 | 0.0858 | -0.41 | 0.30 | -1.35 | 0.3500 |
| AKG, unfiltered & filtered | 0.76 | 0.88 | 0.87 | 0.8070 | -0.35 | 0.20 | -1.74 | 0.2844 |
| AKG, pH adjusted & filtered | -0.79 | 0.75 | -1.05 | 0.6960 | 0.18 | 0.17 | -1.01 | 0.7244 |
| AKG, pH adjusted & unfiltered | -1.56 | 0.88 | -1.78 | 0.2630 | 0.17 | 0.20 | 0.88 | 0.8000 |

**C.** ***C. elegans*** **JU775**

| | General linear model | | | | Random effects Cox Proportional Hazard | | | |
|---|---|---|---|---|---|---|---|---|
| Compound | Effect | Std err | *z*-value | *p*-value | Effect | Std err | *z*-value | *p*-value |
| Thio T, filtered & unfiltered | -16.78 | 1.95 | -8.61 | <1E-05 | 3.74 | 0.41 | 9.11 | <1E-06 |
| AKG, unfiltered & pH adjusted | 2.40 | 1.26 | 1.90 | 0.1310 | -0.35 | 0.29 | -1.21 | 0.4290 |
| AKG, unfiltered & filtered | -1.74 | 1.47 | -1.18 | 0.6140 | 0.14 | 0.28 | 0.51 | 0.9528 |
| AKG, pH adjusted & filtered | -5.28 | 1.34 | -3.94 | <0.001 | 0.69 | 0.28 | 2.43 | 0.0651 |

| | | | | | | | | |
|---|---|---|---|---|---|---|---|---|
| AKG, pH adjusted & unfiltered | -3.54 | 1.48 | -2.39 | 0.0721 | 0.55 | 0.30 | 1.80 | 0.2535 |

**D. *C. briggsae* AF16**

| | General linear model | | | | Random effects Cox Proportional Hazard | | | |
|---|---|---|---|---|---|---|---|---|
| Compound | Effect | Std err | *z*-value | *p*-value | Effect | Std err | *z*-value | *p*-value |
| Thio T, filtered & unfiltered | -11.05 | 1.60 | -6.92 | <1E-04 | 5.48 | 0.43 | 12.83 | <1E-05 |
| AKG, unfiltered & pH adjusted | -0.31 | 1.07 | -0.29 | 0.9510 | 0.16 | 0.25 | 0.63 | 0.7940 |
| AKG, unfiltered & filtered | -7.82 | 1.16 | -6.72 | <0.001 | 1.94 | 0.26 | 7.40 | <0.001 |
| AKG, pH adjusted & filtered | -4.89 | 0.98 | -4.97 | <0.001 | 1.23 | 0.22 | 5.64 | <0.001 |
| AKG, pH adjusted & unfiltered | 2.92 | 1.02 | 2.86 | 0.0195 | -0.71 | 0.23 | -3.12 | 0.0086 |

**E. *C. briggsae* HK104**

| | General linear model | | | | Random effects Cox Proportional Hazard | | | |
|---|---|---|---|---|---|---|---|---|
| Compound | Effect | Std err | z-value | p-value | Effect | Std err | z-value | p-value |
| Thio T, filtered & unfiltered | -18.52 | 2.55 | -7.25 | <0.001 | 4.29 | 0.47 | 9.12 | <0.001 |
| AKG, unfiltered & pH adjusted | 3.87 | 1.83 | 2.12 | 0.0815 | -0.64 | 0.45 | -1.42 | 0.3154 |
| AKG, unfiltered & filtered | -1.84 | 2.09 | -0.88 | 0.7988 | 1.16 | 0.48 | 2.44 | 0.0638 |
| AKG, pH adjusted & filtered | -1.75 | 2.07 | -0.85 | 0.8170 | 0.60 | 0.47 | 1.27 | 0.5568 |
| AKG, pH adjusted & unfiltered | 0.09 | 1.77 | 0.05 | 1.0000 | -0.57 | 0.41 | -1.38 | 0.4864 |

**F. *C. briggsae* JU1348**

| | General linear model | | | | Random effects Cox Proportional Hazard | | | |
|---|---|---|---|---|---|---|---|---|
| Compound | Effect | Std err | z-value | p-value | Effect | Std err | z-value | p-value |
| Thio T, filtered & unfiltered | -14.60 | 1.99 | -7.34 | <0.001 | 6.50 | 0.37 | 17.47 | <0.001 |
| AKG, unfiltered & pH adjusted | 5.21 | 1.97 | 2.65 | 0.0201 | -1.36 | 0.43 | -3.20 | 0.0038 |
| AKG, unfiltered & filtered | 7.62 | 3.54 | 2.16 | 0.1221 | -1.43 | 0.71 | -2.02 | 0.1623 |
| AKG, pH adjusted & filtered | 1.17 | 2.71 | 0.43 | 0.9695 | -0.03 | 0.55 | -0.05 | 1.0000 |
| AKG, pH adjusted & unfiltered | -6.45 | 3.17 | -2.03 | 0.1592 | 1.41 | 0.63 | 2.23 | 0.1049 |
