## Supplementary material for "Automated Lifespan Determination Across *Caenorhabditis* Strains and Species Reveals Assay-Specific Effects of Chemical Interventions": Online Resource 13

**Online Resource 13** Summary of ALM lifespan data under compound treatment (AKG and ThT) conditions, and comparison to median lifespan from comparable manual assays

| | | | | ALM | | | | | | | | Manual | |
|---|---|---|---|---|---|---|---|---|---|---|---|---|---|
| **Species** | **Strain** | **Comp** | **Condition** | **Number of deaths** | **Number censored** | **Total obs.** | **Mean lifespan** | **SEM** | **Median lifespan** | **Lower 95% CI** | **Upper 95% CI** | **Median lifespan** | **% diff med LS from manual** |
| ***C. elegans*** | JU775 | AKG | filter | 184 | 0 | 184 | 24.2 | 0.45 | 24.2 | 22.9 | 25.3 | | |
| | | | nofilter | 297 | 0 | 297 | 18.2 | 0.28 | 17.2 | 16.7 | 18.1 | 25 | -31 |
| | | | pH adjusted | 550 | 0 | 550 | 17.3 | 0.20 | 16.9 | 16.5 | 17.2 | | |
| | | CTRL -H2O | filter | 554 | 5 | 559 | 19.4 | 0.35 | 16.0 | 15.6 | 16.4 | | |
| | | | nofilter | 447 | 0 | 447 | 16.8 | 0.23 | 15.9 | 15.4 | 16.6 | 21 | -24 |
| | | | pH adjusted | 514 | 0 | 514 | 18.8 | 0.29 | 17.0 | 16.5 | 17.4 | | |
| | | ThT | filter | 250 | 0 | 250 | 29.0 | 0.46 | 29.1 | 28.4 | 29.8 | | |
| | | | nofilter | 115 | 0 | 115 | 11.5 | 0.33 | 10.3 | 9.9 | 11.4 | 28 | -63 |
| | MY16 | AKG | filter | 134 | 0 | 134 | 18.3 | 0.24 | 18.1 | 17.4 | 18.5 | | |
| | | | nofilter | 182 | 0 | 182 | 17.7 | 0.27 | 17.0 | 16.7 | 17.8 | 25 | -32 |
| | | | pH adjusted | 492 | 0 | 492 | 17.7 | 0.21 | 17.3 | 16.7 | 17.8 | | |
| | | CTRL -H2O | filter | 384 | 2 | 386 | 17.6 | 0.30 | 15.9 | 14.8 | 16.8 | | |
| | | | nofilter | 307 | 0 | 307 | 15.2 | 0.23 | 14.6 | 14.0 | 15.4 | 16 | -8 |
| | | | pH adjusted | 454 | 0 | 454 | 17.2 | 0.28 | 16.5 | 16.0 | 17.0 | | |
| | | ThT | filter | 181 | 2 | 183 | 27.0 | 0.53 | 28.6 | 27.1 | 29.5 | | |
| | | | nofilter | 43 | 0 | 43 | 17.7 | 0.69 | 18.7 | 17.3 | 19.4 | 28 | -33 |
| | N2 | AKG | filter | 181 | 0 | 181 | 19.8 | 0.24 | 19.3 | 18.7 | 20.1 | | |
| | | | nofilter | 327 | 1 | 328 | 14.1 | 0.11 | 14.0 | 13.9 | 14.3 | 23 | -39 |
| | | | pH adjusted | 560 | 0 | 560 | 16.8 | 0.14 | 16.7 | 16.3 | 17.1 | | |
| | | CTRL -H2O | filter | 495 | 1 | 496 | 18.0 | 0.23 | 18.0 | 17.1 | 18.5 | | |
| | | | nofilter | 513 | 0 | 513 | 15.9 | 0.13 | 16.5 | 16.2 | 16.8 | 18 | -8 |
| | | | pH adjusted | 660 | 0 | 660 | 17.0 | 0.13 | 17.5 | 17.2 | 17.8 | | |
| | | ThT | filter | 392 | 3 | 395 | 23.9 | 0.36 | 24.9 | 23.9 | 25.6 | | |
| | | | nofilter | 148 | 0 | 148 | 9.8 | 0.16 | 9.4 | 8.9 | 9.7 | 23 | -59 |
| ***C. briggsae*** | AF16 | AKG | filter | 75 | 0 | 75 | 26.5 | 0.62 | 27.1 | 24.4 | 28.6 | | |
| | | | nofilter | 163 | 0 | 163 | 17.1 | 0.32 | 16.6 | 16.2 | 17.5 | 28 | -41 |
| | | | pH adjusted | 325 | 0 | 325 | 17.0 | 0.21 | 16.7 | 16.2 | 17.1 | | |
| | | CTRL -H2O | filter | 213 | 1 | 214 | 23.0 | 0.43 | 23.2 | 22.3 | 24.5 | | |
| | | | nofilter | 325 | 2 | 327 | 20.8 | 0.25 | 21.1 | 20.6 | 21.7 | 26 | -19 |
| | | | pH adjusted | 392 | 0 | 392 | 19.9 | 0.28 | 19.4 | 18.6 | 20.3 | | |
| | | ThT | filter | 70 | 0 | 70 | 21.3 | 1.21 | 19.6 | 14.4 | 24.1 | | |
| | | | nofilter | 109 | 0 | 109 | 9.1 | 0.06 | 9.1 | 9.0 | 9.2 | 32 | -72 |
| | HK104 | AKG | filter | 64 | 0 | 64 | 34.3 | 1.13 | 36.8 | 33.8 | 39.3 | | |
| | | | nofilter | 318 | 5 | 323 | 23.5 | 0.39 | 22.5 | 21.5 | 22.9 | 35 | -36 |
| | | | pH adjusted | 406 | 0 | 406 | 23.6 | 0.31 | 24.3 | 23.1 | 25.0 | | |
| | | CTRL -H2O | filter | 282 | 1 | 283 | 35.1 | 0.60 | 36.8 | 35.2 | 37.7 | | |
| | | | nofilter | 488 | 5 | 493 | 26.5 | 0.30 | 26.5 | 25.8 | 27.2 | 38 | -30 |
| | | | pH adjusted | 553 | 0 | 553 | 30.6 | 0.28 | 31.5 | 30.9 | 32.1 | | |
| | | ThT | filter | 146 | 2 | 148 | 40.4 | 1.13 | 44.1 | 42.0 | 45.8 | | |
| | | | nofilter | 93 | 1 | 94 | 13.6 | 0.47 | 12.0 | 11.6 | 12.3 | 42 | -71 |
| | JU1348 | AKG | filter | 57 | 0 | 57 | 20.4 | 0.59 | 19.1 | 18.7 | 20.0 | | |
| | | | nofilter | 105 | 0 | 105 | 20.9 | 0.46 | 20.4 | 19.6 | 21.2 | 25 | -18 |
| | | | pH adjusted | 432 | 0 | 432 | 16.4 | 0.22 | 15.7 | 15.2 | 16.2 | | |
| | | CTRL -H2O | filter | 208 | 0 | 208 | 24.1 | 0.57 | 23.0 | 21.6 | 24.1 | | |
| | | | nofilter | 279 | 0 | 279 | 19.2 | 0.36 | 18.0 | 17.3 | 18.4 | 26 | -31 |
| | | | pH adjusted | 337 | 0 | 337 | 21.1 | 0.30 | 20.4 | 19.8 | 21.5 | | |
| | | ThT | filter | 172 | 0 | 172 | 26.2 | 0.46 | 25.7 | 24.6 | 26.5 | | |
| | | | nofilter | 72 | 0 | 72 | 10.0 | 0.14 | 9.8 | 9.2 | 10.6 | 28 | -65 |
