## Supplementary material for "Automated Lifespan Determination Across *Caenorhabditis* Strains and Species Reveals Assay-Specific Effects of Chemical Interventions": Online Resource 14

**Online Resource 14** Variance components estimates for longevity for the NP1, PG, and RVL compound experiments, analyzed separately for each strain. Values are from a hierarchical randomized block design estimated either via a restricted maximum likelihood general linear model using the *lme4* package (v. 1.1-21) or via a random effects Cox Proportional Hazards model as implemented by the *coxme* package (v. 2.2-10) in R (Therneau 2012).

**A.** ***C. elegans*** **N2** (*n* = 1,806)

| | General Linear Model | | | | Cox Prop Hazard |
|---|---|---|---|---|---|
| **Source** | **Var Comp** | **Lower 95% CI** | **Upper 95% CI** | **Percent Total** | **Var Comp** |
| Lab | 1.09 | 0.00 | 8.32 | 9.9 | 0.08 |
| Scanner[Lab] | 0.00 | 0.00 | 3.10 | 0.0 | 0.01 |
| Trial[Lab,Scn] | 1.55 | 0.18 | 3.94 | 14.1 | 0.39 |
| Plate-T[Lab,Scn,Trial] | 0.33 | 0.08 | 0.67 | 3.0 | 0.06 |
| Residual | 8.01 | 7.51 | 8.57 | 72.9 | |
| Total | 10.99 | | | 100.0 | |

**B.** ***C. elegans*** **MY16** (*n* = 1,539)

| | General Linear Model | | | | Cox Prop Hazard |
|---|---|---|---|---|---|
| **Source** | **Var Comp** | **Lower 95% CI** | **Upper 95% CI** | **Percent Total** | **Var Comp** |
| Lab | 0.15 | 0.00 | 4.46 | 1.0 | 0.00 |
| Scanner[Lab] | 0.00 | 0.00 | 2.78 | 0.0 | 0.01 |
| Trial[Lab,Scn] | 2.15 | 0.69 | 5.53 | 13.7 | 0.12 |
| Plate-T[Lab,Scn,Trial] | 2.43 | 1.28 | 3.89 | 15.6 | 0.26 |
| Residual | 10.92 | 10.17 | 11.74 | 69.8 | |
| Total | 15.65 | | | 100.0 | |

**C.** ***C. elegans*** **JU775** (*n* = 1,695)

| | General Linear Model | | | | Cox Prop Hazard |
|---|---|---|---|---|---|
| **Source** | **Var Comp** | **Lower 95% CI** | **Upper 95% CI** | **Percent Total** | **Var Comp** |
| Lab | 0.00 | 0.00 | 1.78 | 0.0 | 0.02 |
| Scanner[Lab] | 0.00 | 0.00 | 1.50 | 0.0 | 0.00 |
| Trial[Lab,Scn] | 1.95 | 0.77 | 4.45 | 8.7 | 0.11 |
| Plate-T[Lab,Scn,Trial] | 0.72 | 0.15 | 1.45 | 3.2 | 0.07 |
| Residual | 19.79 | 18.50 | 21.20 | 88.1 | |
| Total | 22.46 | | | 100.0 | |

**D. *C. briggsae* AF16** (*n* = 1,029)

| Source | General Linear Model | | | | Cox Prop Hazard |
|---|---|---|---|---|---|
| | Var Comp | Lower 95% CI | Upper 95% CI | Percent Total | Var Comp |
| Lab | 0.00 | 0.00 | 2.98 | 0.0 | 0.00 |
| Scanner[Lab] | 0.00 | 0.00 | 2.99 | 0.0 | 0.00 |
| Trial[Lab,Scn] | 2.81 | 0.63 | 7.64 | 9.0 | 0.17 |
| Plate-T[Lab,Scn,Trial] | 3.12 | 1.29 | 5.39 | 10.0 | 0.10 |
| Residual | 25.13 | 23.03 | 27.49 | 80.9 | |
| Total | 31.06 | | | 100.0 | |

**E. *C. briggsae* HK104** (*n* = 1,535)

| Source | General Linear Model | | | | Cox Prop Hazard |
|---|---|---|---|---|---|
| | Var Comp | Lower 95% CI | Upper 95% CI | Percent Total | Var Comp |
| Lab | 17.35 | 2.99 | 103.42 | 31.4 | 0.47 |
| Scanner[Lab] | 0.79 | 0.00 | 4.22 | 1.4 | 0.07 |
| Trial[Lab,Scn] | 0.00 | 0.00 | 3.19 | 0.0 | 0.00 |
| Plate-T[Lab,Scn,Trial] | 2.57 | 1.10 | 4.75 | 4.7 | 0.14 |
| Residual | 34.54 | 32.18 | 37.14 | 62.5 | |
| Total | 55.24 | | | 100.0 | |

**F. *C. briggsae* JU1348** (*n* = 1,242)

| Source | General Linear Model | | | | Cox Prop Hazard |
|---|---|---|---|---|---|
| | Var Comp | Lower 95% CI | Upper 95% CI | Percent Total | Var Comp |
| Lab | 4.55 | 0.00 | 31.24 | 14.8 | 0.19 |
| Scanner[Lab] | 2.01 | 0.64 | 7.02 | 6.5 | 0.14 |
| Trial[Lab,Scn] | 0.00 | 0.00 | 4.24 | 0.0 | 0.00 |
| Plate-T[Lab,Scn,Trial] | 1.29 | 0.42 | 2.39 | 4.2 | 0.07 |
| Residual | 22.85 | 21.11 | 24.77 | 74.4 | |
| Total | 30.70 | | | 100.0 | |

**Reference List**

Therneau T., 2012 Mixed Effects Cox Models R Foundation for Statistical Computing R package
