## Supplementary material for "Automated Lifespan Determination Across *Caenorhabditis* Strains and Species Reveals Assay-Specific Effects of Chemical Interventions": Online Resource 15

**Online Resource 15** Variance components estimates for longevity for the Thioflavin T filtered and unfiltered compound experiments, analyzed separately for each strain. Values are from a hierarchical randomized block design estimated either via a restricted maximum likelihood general linear model using the *lme4* package (v. 1.1-21) or via a random effects Cox Proportional Hazards model as implemented by the *coxme* package (v. 2.2-10) in R (Therneau 2012).

**A. *C. elegans* N2** (*n* = 1,552)

| Source | General Linear Model | | | | Cox Prop Hazard |
|---|---|---|---|---|---|
| | Var Comp | Lower 95% CI | Upper 95% CI | Percent Total | Var Comp |
| Lab | 0.33 | 0.00 | 3.14 | 1.29 | 0.00 |
| Scanner[Lab] | 0.00 | 0.00 | 2.12 | 0.00 | 0.02 |
| Trial[Lab,Scn] | 0.18 | 0.00 | 2.00 | 0.72 | 0.02 |
| Plate-T[Lab,Scn,Trial] | 2.90 | 1.08 | 4.93 | 11.54 | 0.14 |
| Residual | 21.76 | 20.27 | 23.39 | 86.45 | |
| Total | 25.17 | | | 100.0 | |

**B. *C. elegans* MY16** (*n* = 919)

| Source | General Linear Model | | | | Cox Prop Hazard |
|---|---|---|---|---|---|
| | Var Comp | Lower 95% CI | Upper 95% CI | Percent Total | Var Comp |
| Lab | 0.00 | 0.00 | 4.59 | 0.00 | 0.00 |
| Scanner[Lab] | 2.01 | 0.00 | 12.46 | 5.99 | 0.00 |
| Trial[Lab,Scn] | 4.33 | 1.36 | 13.08 | 12.94 | 0.22 |
| Plate-T[Lab,Scn,Trial] | 2.81 | 0.49 | 6.39 | 8.41 | 0.05 |
| Residual | 24.31 | 22.18 | 26.73 | 72.66 | |
| Total | 33.45 | | | 100.0 | |

**C. *C. elegans* JU775** (*n* = 1,371)

| Source | General Linear Model | | | | Cox Prop Hazard |
|---|---|---|---|---|---|
| | Var Comp | Lower 95% CI | Upper 95% CI | Percent Total | Var Comp |
| Lab | 0.71 | 0.00 | 8.44 | 1.57 | 0.00 |
| Scanner[Lab] | 3.23 | 0.00 | 9.58 | 7.12 | 0.12 |
| Trial[Lab,Scn] | 0.00 | 0.00 | 6.19 | 0.00 | 0.00 |
| Plate-T[Lab,Scn,Trial] | 6.04 | 2.57 | 10.49 | 13.32 | 0.29 |
| Residual | 35.34 | 32.77 | 38.16 | 77.99 | |
| Total | 45.31 | | | 100.0 | |

**D.** ***C. briggsae*** **AF16** (*n* = 720)

| Source | General Linear Model | | | | Cox Prop Hazard |
|---|---|---|---|---|---|
| | Var Comp | Lower 95% CI | Upper 95% CI | Percent Total | Var Comp |
| Lab | 2.68 | 0.00 | 21.07 | 7.64 | 0.31 |
| Scanner[Lab] | 0.00 | 0.00 | 3.84 | 0.00 | 0.00 |
| Trial[Lab,Scn] | 5.70 | 0.00 | 13.58 | 16.23 | 0.47 |
| Plate-T[Lab,Scn,Trial] | 1.91 | 0.00 | 17.51 | 5.43 | 0.08 |
| Residual | 24.83 | 22.37 | 27.68 | 70.70 | |
| Total | 35.12 | | | 100.0 | |

**E.** ***C. briggsae*** **HK104** (*n* = 1,018)

| Source | General Linear Model | | | | Cox Prop Hazard |
|---|---|---|---|---|---|
| | Var Comp | Lower 95% CI | Upper 95% CI | Percent Total | Var Comp |
| Lab | 41.15 | 0.00 | 13.08 | 35.94 | 0.85 |
| Scanner[Lab] | 16.43 | 4.45 | 45.17 | 14.35 | 1.07 |
| Trial[Lab,Scn] | 0.00 | 0.00 | 10.94 | 0.00 | 0.00 |
| Plate-T[Lab,Scn,Trial] | 7.04 | 2.75 | 13.08 | 6.15 | 0.23 |
| Residual | 49.86 | 45.69 | 54.55 | 43.55 | |
| Total | 114.49 | | | 100.0 | |

**F.** ***C. briggsae*** **JU1348** (*n* = 731)

| Source | General Linear Model | | | | Cox Prop Hazard |
|---|---|---|---|---|---|
| | Var Comp | Lower 95% CI | Upper 95% CI | Percent Total | Var Comp |
| Lab | 3.35 | 0.00 | 28.55 | 6.87 | 0.03 |
| Scanner[Lab] | 8.29 | 0.00 | 24.56 | 17.01 | 0.93 |
| Trial[Lab,Scn] | 0.00 | 0.00 | 8.70 | 0.00 | 0.00 |
| Plate-T[Lab,Scn,Trial] | 3.23 | 0.46 | 7.24 | 6.62 | 0.01 |
| Residual | 33.88 | 30.58 | 37.72 | 69.50 | |
| Total | 48.74 | | | 100.0 | |

### Reference List

Therneau T., 2012 Mixed Effects Cox Models R Foundation for Statistical Computing R package
