## Supplementary material for "Automated Lifespan Determination Across *Caenorhabditis* Strains and Species Reveals Assay-Specific Effects of Chemical Interventions": Online Resource 16

**Online Resource 16** Variance components estimates for longevity for the α-ketoglutarate pH adjusted and unfiltered compound experiments, analyzed separately for each strain. Values are from a hierarchical randomized block design estimated either via a restricted maximum likelihood general linear model using the *lme4* package (v. 1.1-21) or via a random effects Cox Proportional Hazards model as implemented by the *coxme* package (v. 2.2-10) in R (Therneau 2012).

**A.** ***C. elegans*** **N2** (*n* = 2,061)

| | General Linear Model | | | | Cox Prop Hazard |
|---|---|---|---|---|---|
| **Source** | **Var Comp** | **Lower 95% CI** | **Upper 95% CI** | **Percent Total** | **Var Comp** |
| Lab | 0.75 | 0.00 | 5.80 | 7.35 | 0.20 |
| Scanner[Lab] | 0.00 | 0.00 | 2.65 | 0.00 | 0.00 |
| Trial[Lab,Scn] | 2.02 | 0.01 | 4.32 | 19.79 | 0.20 |
| Plate-T[Lab,Scn,Trial] | 1.31 | 0.63 | 2.27 | 12.86 | 0.35 |
| Residual | 6.13 | 5.77 | 6.53 | 60.00 | |
| Total | 10.22 | | | 100.0 | |

**B.** ***C. elegans*** **MY16** (*n* = 1,435)

| | General Linear Model | | | | Cox Prop Hazard |
|---|---|---|---|---|---|
| **Source** | **Var Comp** | **Lower 95% CI** | **Upper 95% CI** | **Percent Total** | **Var Comp** |
| Lab | 0.00 | 0.00 | 2.46 | 0.00 | 0.00 |
| Scanner[Lab] | 2.67 | 0.00 | 7.61 | 11.17 | 0.23 |
| Trial[Lab,Scn] | 1.07 | 0.00 | 10.87 | 4.47 | 0.01 |
| Plate-T[Lab,Scn,Trial] | 1.63 | 0.58 | 3.43 | 6.84 | 0.16 |
| Residual | 18.51 | 17.21 | 19.96 | 77.53 | |
| Total | 23.87 | | | 100.0 | |

**C.** ***C. elegans*** **JU775** (*n* = 1,808)

| | General Linear Model | | | | Cox Prop Hazard |
|---|---|---|---|---|---|
| **Source** | **Var Comp** | **Lower 95% CI** | **Upper 95% CI** | **Percent Total** | **Var Comp** |
| Lab | 0.43 | | | | 0.04 |
| Scanner[Lab] | 0.00 | | | | 0.22 |
| Trial[Lab,Scn] | 4.38 | | | | 0.04 |
| Plate-T[Lab,Scn,Trial] | 3.70 | | | | 0.18 |
| Residual | 19.25 | | | | |
| Total | 27.76 | | | | |

**D.** ***C. briggsae*** **AF16** (*n* = 1,207)

| Source | General Linear Model | | | | Cox Prop Hazard |
|---|---|---|---|---|---|
| | Var Comp | Lower 95% CI | Upper 95% CI | Percent Total | Var Comp |
| Lab | 0.29 | 0.00 | 3.98 | 1.30 | 0.00 |
| Scanner[Lab] | 2.04 | 0.00 | 5.77 | 9.09 | 0.12 |
| Trial[Lab,Scn] | 0.02 | 0.00 | 8.08 | 0.08 | 0.00 |
| Plate-T[Lab,Scn,Trial] | 2.33 | 0.47 | 4.28 | 10.39 | 0.12 |
| Residual | 17.73 | 16.37 | 19.28 | 79.15 | |
| Total | 22.40 | | | 100.0 | |

**E.** ***C. briggsae*** **HK104** (*n* = 1,775)

| Source | General Linear Model | | | | Cox Prop Hazard |
|---|---|---|---|---|---|
| | Var Comp | Lower 95% CI | Upper 95% CI | Percent Total | Var Comp |
| Lab | 0.01 | 0.00 | 14.15 | 0.02 | 0.01 |
| Scanner[Lab] | 7.50 | 1.90 | 13.59 | 15.84 | 0.31 |
| Trial[Lab,Scn] | 0.00 | 0.00 | 18.63 | 0.00 | 0.01 |
| Plate-T[Lab,Scn,Trial] | 8.65 | 4.63 | 11.38 | 18.27 | 0.53 |
| Residual | 31.19 | 29.19 | 33.37 | 65.87 | |
| Total | 47.34 | | | 100.0 | |

**F.** ***C. briggsae*** **JU1348** (*n* = 1,153)

| Source | General Linear Model | | | | Cox Prop Hazard |
|---|---|---|---|---|---|
| | Var Comp | Lower 95% CI | Upper 95% CI | Percent Total | Var Comp |
| Lab | 0.79 | 0.00 | 7.19 | 2.67 | 0.00 |
| Scanner[Lab] | 0.00 | 0.00 | 4.93 | 0.00 | 0.00 |
| Trial[Lab,Scn] | 0.00 | 0.00 | 4.68 | 0.00 | 0.00 |
| Plate-T[Lab,Scn,Trial] | 7.37 | 3.29 | 11.57 | 24.94 | 0.36 |
| Residual | 21.39 | 19.71 | 23.28 | 72.39 | |
| Total | 29.55 | | | 100.0 | |

**Reference List**

Therneau T., 2012 Mixed Effects Cox Models R Foundation for Statistical Computing R package
