## Supplementary material for "Automated Lifespan Determination Across *Caenorhabditis* Strains and Species Reveals Assay-Specific Effects of Chemical Interventions": Online Resource 17

**Online Resource 17** Variance components estimates for longevity for the α-ketoglutarate pH adjusted, filtered, and unfiltered compound experiments from the Phillips laboratory, analyzed separately for each strain. Values are from a hierarchical randomized block design estimated either via a restricted maximum likelihood general linear model using the *lme4* package (v. 1.1-21) or via a random effects Cox Proportional Hazards model as implemented by the *coxme* package (v. 2.2-10) in R (Therneau 2012).

**A.** ***C. elegans*** **N2** (*n* = 1,436)

| | **General Linear Model** | | | | **Cox Prop Hazard** |
|---|---|---|---|---|---|
| **Source** | **Var Comp** | **Lower 95% CI** | **Upper 95% CI** | **Percent Total** | **Var Comp** |
| Scanner | 0.00 | 0.00 | 0.42 | 0.00 | 0.00 |
| Trial*Scanner | 0.00 | 0.00 | 0.41 | 0.00 | 0.00 |
| Plate-T[Scanner,Trial] | 1.54 | 0.71 | 2.21 | 13.68 | 0.11 |
| Residual | 9.69 | 9.01 | 10.45 | 86.32 | |
| Total | 11.23 | | | 100.0 | |

**B.** ***C. elegans*** **MY16** (*n* = 856)

| | **General Linear Model** | | | | **Cox Prop Hazard** |
|---|---|---|---|---|---|
| **Source** | **Var Comp** | **Lower 95% CI** | **Upper 95% CI** | **Percent Total** | **Var Comp** |
| Scanner[Lab] | 0.49 | 0.00 | 2.33 | 2.30 | 0.00 |
| Trial[Lab,Scn] | 0.92 | 0.00 | 2.73 | 4.33 | 0.07 |
| Plate-T[Lab,Scn,Trial] | 0.00 | 0.00 | 0.69 | 0.00 | 0.00 |
| Residual | 19.78 | 17.95 | 21.75 | 93.37 | |
| Total | 21.19 | | | 100.0 | |

**C.** ***C. elegans*** **JU775** (*n* = 1,372)

| | **General Linear Model** | | | | **Cox Prop Hazard** |
|---|---|---|---|---|---|
| **Source** | **Var Comp** | **Lower 95% CI** | **Upper 95% CI** | **Percent Total** | **Var Comp** |
| Scanner[Lab] | 0.00 | 0.00 | 3.47 | 0.00 | 0.00 |
| Trial[Lab,Scn] | 1.97 | 0.00 | 4.65 | 4.93 | 0.00 |
| Plate-T[Lab,Scn,Trial] | 1.53 | 0.03 | 3.85 | 3.82 | 0.09 |
| Residual | 36.54 | 33.91 | 39.47 | 91.26 | |
| Total | 40.04 | | | 100.0 | |

**D.** ***C. briggsae*** **AF16** (*n* = 744)

| Source | General Linear Model | | | | Cox Prop Hazard |
|---|---|---|---|---|---|
| | Var Comp | Lower 95% CI | Upper 95% CI | Percent Total | Var Comp |
| Scanner[Lab] | 0.00 | 0.00 | 3.35 | 0.00 | 0.00 |
| Trial[Lab,Scn] | 3.93 | 0.81 | 6.98 | 15.66 | 0.16 |
| Plate-T[Lab,Scn,Trial] | 0.00 | 0.00 | 0.78 | 0.00 | 0.00 |
| Residual | 21.17 | 19.06 | 23.43 | 84.34 | |
| Total | 25.10 | | | 100.0 | |

**E.** ***C. briggsae*** **HK104** (*n* = 1,179)

| Source | General Linear Model | | | | Cox Prop Hazard |
|---|---|---|---|---|---|
| | Var Comp | Lower 95% CI | Upper 95% CI | Percent Total | Var Comp |
| Scanner[Lab] | 0.87 | 1.18 | 6.26 | 2.08 | 0.08 |
| Trial[Lab,Scn] | 0.01 | 0.00 | 4.35 | 0.01 | 0.01 |
| Plate-T[Lab,Scn,Trial] | 3.59 | 0.00 | 3.65 | 8.63 | 0.21 |
| Residual | 37.17 | 34.28 | 40.38 | 89.28 | |
| Total | 41.63 | | | 100.0 | |

**F.** ***C. briggsae*** **JU1348** (*n* = 602)

| Source | General Linear Model | | | | Cox Prop Hazard |
|---|---|---|---|---|---|
| | Var Comp | Lower 95% CI | Upper 95% CI | Percent Total | Var Comp |
| Scanner[Lab] | 0.00 | 0.00 | 3.46 | 0.00 | 0.00 |
| Trial[Lab,Scn] | 0.00 | 0.00 | 4.32 | 0.00 | 0.00 |
| Plate-T[Lab,Scn,Trial] | 5.13 | 1.26 | 7.80 | 13.17 | 0.22 |
| Residual | 33.80 | 30.42 | 37.71 | 86.83 | |
| Total | 38.93 | | | 100.0 | |

**Reference List**

Therneau T., 2012 Mixed Effects Cox Models R Foundation for Statistical Computing R package
