## Supplementary material for "Automated Lifespan Determination Across *Caenorhabditis* Strains and Species Reveals Assay-Specific Effects of Chemical Interventions": Online Resource 18

**Online Resource 18 Thioflavin T is toxic under the intense illumination during automated lifespan analysis**

Survivorship curves generated using automated lifespan analysis under control (black lines) or thioflavin T (mustard lines) under normal illumination (solid lines) or filtered (dash-dot lines) Vertical dotted line shows day plates introduced to the scanners and the start of intense illumination. Automated analysis resulted in shortened lifespan under thioflavin T exposure, while filtering the ALM light restored the lifespan extension of thioflavin T exposure.

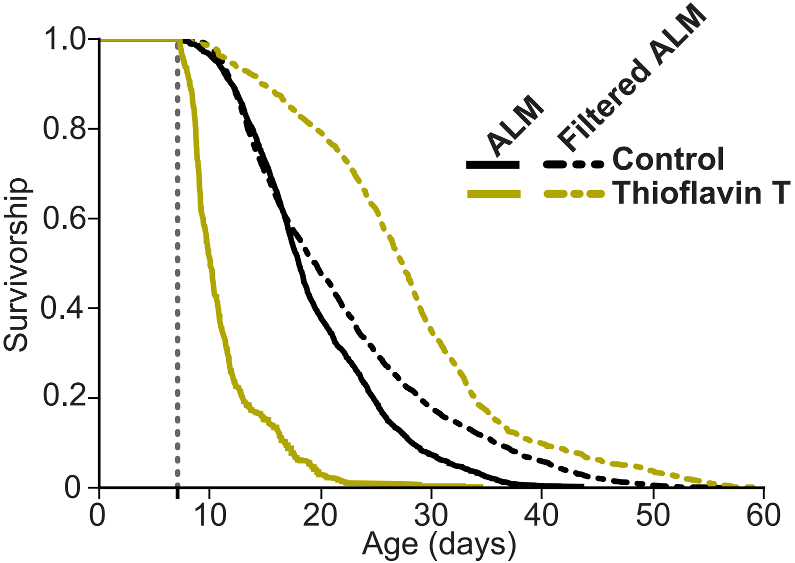

Survivorship
1.0
0.8
0.6
0.4
0.2
0
0
10
20
30
40
50
60
Age (days)
ALM
Filtered ALM
Control
Thioflavin T
