## Supplementary material for "Automated Lifespan Determination Across *Caenorhabditis* Strains and Species Reveals Assay-Specific Effects of Chemical Interventions": Online Resource 19

**Online Resource 19 α-ketoglutarate lifespan effects are not due to pH differences in automated lifespan analysis**

The percent change in median lifespan from control for animals grown under adult exposure to α-ketoglutarate for three *C. elegans* (N2, JU775, and MY16) and *C. briggsae* (AF16, JU1348, and HK104) strains. Each point represents the change in median lifespan for an individual plate trial. Replicates were generated at three CITP sites (Blue-Buck Institute, Green-Oregon and Red- Rutgers). Lifespans were measured by standard manual lifespan analysis (open circles), ALM analysis (closed circles), or ALM analysis in which the α-ketoglutarate stock solutions had been adjusted to pH 6 (closed squares) prior to plate treatment (see materials and methods). Asterisks represent *p*-values (****$p<0.0001$, *** $p<0.001$, ** $p<0.01$ and * $p<0.05$) from the CPH model when comparing the lifespans under compound exposure versus the lifespans exposed to the vehicle control.

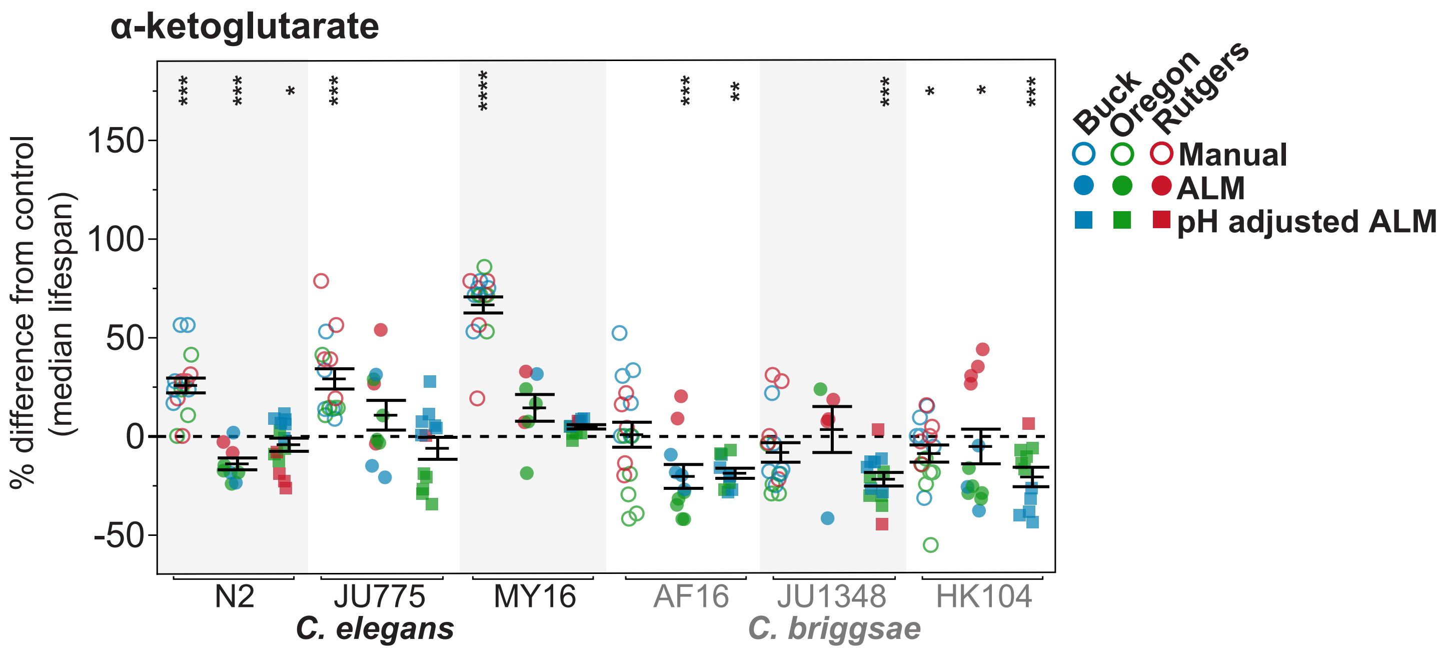

α-ketoglutarate
% difference from control (median lifespan)
150
100
50
0
-50
***
***
*
***
****
***
**
***
*
*
***
N2
JU775
MY16
AF16
JU1348
HK104
C. elegans
C. briggsae
Buck
Oregon
Rutgers
Manual
ALM
pH adjusted ALM
