## Supplementary material for "Automated Lifespan Determination Across *Caenorhabditis* Strains and Species Reveals Assay-Specific Effects of Chemical Interventions": Online Resource 20

**Online Resource 20** Manual vs. ALM: approximate average return rates per plate (observed mean number of deaths versus expected number of animals at experiment start). The expected number of animals per plate for manual assays was 37.5, and for ALM assays was 50.

| | | Manual | | ALM | |
|---|---|---|---|---|---|
| | **Strain** | **Observed deaths** | **% expected observed** | **Observed deaths** | **% expected observed** |
| *C. elegans* | N2 | 31 | 83 | 38 | 76 |
| | N2 PD1073 | - | - | 33 | 66 |
| | CB4856 | 31 | 83 | 30 | 60 |
| | ED3040 | 33 | 88 | 37 | 74 |
| | JU775 | 32 | 85 | 34 | 68 |
| | JU1088 | 33 | 88 | 33 | 66 |
| | JU1652 | 32 | 85 | 32 | 64 |
| | MY16 | 31 | 83 | 30 | 60 |
| | QX1211 | 31 | 83 | 26 | 52 |
| *C. briggsae* | AF16 | 25 | 67 | 20 | 40 |
| | ED3092 | 25 | 67 | 29 | 58 |
| | HK104 | 25 | 67 | 29 | 58 |
| | JU726 | 29 | 77 | 24 | 48 |
| | JU1264 | 24 | 64 | 29 | 58 |
| | JU1348 | 24 | 64 | 24 | 48 |
| | NIC20 | 25 | 67 | 29 | 58 |
| | QR25 | 25 | 67 | 32 | 64 |
| *C. tropicalis* | JU1373 | 30 | 80 | 33 | 66 |
| | JU1630 | 31 | 83 | 20 | 40 |
| | NIC58 | 25 | 67 | 23 | 46 |
| | NIC122 | 23 | 61 | 27 | 54 |
| | QG131 | 25 | 67 | 27 | 54 |
| | QG834 | 29 | 77 | 29 | 58 |
